# Nanoscale imaging of bacterial infections by sphingolipid expansion microscopy

**DOI:** 10.1101/2020.05.06.080663

**Authors:** Ralph Götz, Tobias C. Kunz, Julian Fink, Franziska Solger, Jan Schlegel, Jürgen Seibel, Vera Kozjak-Pavlovic, Thomas Rudel, Markus Sauer

**Author notes:** Corresponding authors, (T.R.), (M.S.). These authors contributed equally: Ralph Götz, Tobias C. Kunz.

## Abstract

Expansion microscopy (ExM) enables super-resolution imaging of proteins and nucleic acids on conventional microscopes. However, imaging of details of the organization of lipid bilayers by light microscopy remains challenging. We introduce an azide- and amino-modified sphingolipid ceramide, which upon incorporation into membranes can be labeled by click chemistry and linked into hydrogels, followed by 4x to 10x expansion. Confocal and structured illumination microscopy (SIM) enabled imaging of sphingolipids and their interactions with proteins in the membrane of intracellular organelles with a spatial resolution of 10-20 nm. Because sphingolipids accumulated efficiently in pathogens we used sphingolipid ExM to investigate bacterial infections of human HeLa229 cells by *Neisseria gonorrhoeae, Chlamydia trachomatis* and *Simkania negevensis* with a resolution so far only provided by electron microscopy. In particular, sphingolipid ExM allowed us to visualize the inner and outer membrane of intracellular bacteria and determine their distance to 27.6 ± 7.7 nm.

## Introduction

In the last decade, super-resolution microscopy has evolved as a very powerful method for subdiffraction-resolution fluorescence imaging of cells and structural investigations of cellular organelles.^1,2^ Super-resolution microscopy methods can now provide a spatial resolution that is well below the diffraction limit of light microscopy, enabling invaluable insights into the spatial organization of proteins in biological samples. However, in particular three-dimensional and multicolor super-resolution microscopy methods require elaborate equipment and experience and are therefore mostly restricted to specialized laboratories.

Expansion microscopy (ExM) provides an alternative approach to bypass the diffraction limit and enable super-resolution imaging on standard fluorescence microscopes. By linking a protein of interest into a dense, cross-linked network of a swellable polyelectrolyte hydrogel, biological specimens can be physically expanded allowing ∼70 nm lateral resolution by confocal laser scanning microscopy. Since its introduction by Boyden and co-workers in 2015^3^, expansion microscopy (ExM) has shown impressive results including the magnified visualization of pre- or post-expansion labeled proteins and RNAs with fluorescent proteins, antibodies, and oligonucleotides, respectively, in cells, tissues, and human clinical specimen.^4^ ExM has been developing at an enormous speed with various protocols providing expansion factors from 4x^3^ to 10x^5,6^ and even 20x by iterative expansion.^7^ In addition, various protocols have been introduced enabling subdiffraction-resolution imaging of proteins, RNA, and bacteria in cultured cells, neurons, and tissues by confocal fluorescence microscopy and in combination with super-resolution microscopy.^5-14^

In order to be usable for ExM, the molecule of interest has to exhibit amino groups that can react with glutaraldehyde (GA)^9^, MA-NHS^9^, AcX^10^, or Label-X^11^ and be linked into the polyelectrolyte hydrogel. The plasma membrane of cells is mainly composed of glycerophospholipids, sphingolipids, and cholesterol. Due to the lack of primary amino groups, these lipids neither can be fixed by formaldehyde, glutaraldehyde and other chemical fixatives nor expanded using available ExM protocols. To this end, we sought to functionalize a lipid that is compatible with ExM. So far, sphingolipids have only been functionalized as azides to enable fluorescence labeling by click chemistry after incorporation into cellular membranes.^15-18^ Therefore, we set out to introduce an azide and primary amino group into sphingolipids to enable fluorescence labeling and chemical fixation as well as linking of the lipid into a swellable hydrogel. Our results demonstrate that the designed bifunctional sphingolipid is efficiently incorporated into membranes of cells and especially in bacterial membranes, which allowed us to investigate the distribution of lipids and interactions with proteins in cellular and bacterial membranes with high spatial resolution.

## Results and Discussion

### Sphingolipid ExM of cellular membranes

Sphingolipids are natural lipids comprised of the sphingoid base backbone sphingosine, which when N-acylated with fatty acids forms ceramide, a central molecule in sphingolipid biology. Sphingolipid ceramides regulate cellular processes such as differentiation, proliferation, growth arrest and apoptosis. Ceramide-rich membrane areas promote structural changes within the plasma membrane, which segregate membrane receptors and affect the membrane curvature and vesicle formation, fusion and trafficking.^19,20^

We selected ω-N_3_-C_6_-ceramide, which is efficiently incorporated into cellular membranes and can be click-labeled with DBCO-functionalized dyes for fluorescence imaging^21,22^, for further functionalization with a primary amino group (Supporting Information and Supplementary Figures S1-S13). Therefore, we synthesized α-NH_2_-ω-N_3_-C_6_-ceramide from (*tert*-butoxycarbonyl)-L-lysine (Figure 1A). We first assessed if the synthesized α-NH_2_-ω-N_3_-C_6_-ceramide (Figure 1A) is incorporated into cellular membranes similar to the control ceramide without amino modification and can be labeled by click chemistry with DBCO-dyes. For this, cells were fed for 1 h with the two ceramides, fixed with glutaraldehyde and click-labeled with DBCO-Alexa Fluor 488. Confocal fluorescence images showed that both analogues ω-N_3_-C_6_-ceramide and α-NH_2_-ω-N_3_-C_6_-ceramide were incorporated into the plasma membrane and membranes of intracellular organelles of HeLa229 cells with comparable efficiency (Figure 1). Fluorescence recovery after photobleaching (FRAP) experiments with both ceramides indicated that ω-N_3_-C_6_-ceramide shows a higher mobility in the plasma membrane after fixation than α-NH_2_-ω-N_3_-C_6_-ceramide (Figure 1B). This finding was corroborated by the treatment of labeled cells with detergents, which wash out unfixed lipids. Upon addition of Triton X-100 or saponine ω-N_3_-C_6_-ceramide was efficiently washed out whereas the fluorescence signal of the amino-modified analog α-NH_2_-ω-N_3_-C_6_-ceramide decreased only slightly and was preserved for weeks (Figure 1C and Supplementary Figure S14). These results demonstrate that the crosslinker glutaraldehyde can efficiently fix amino-modified ceramides incorporated into cellular membranes.

**Figure 1.**
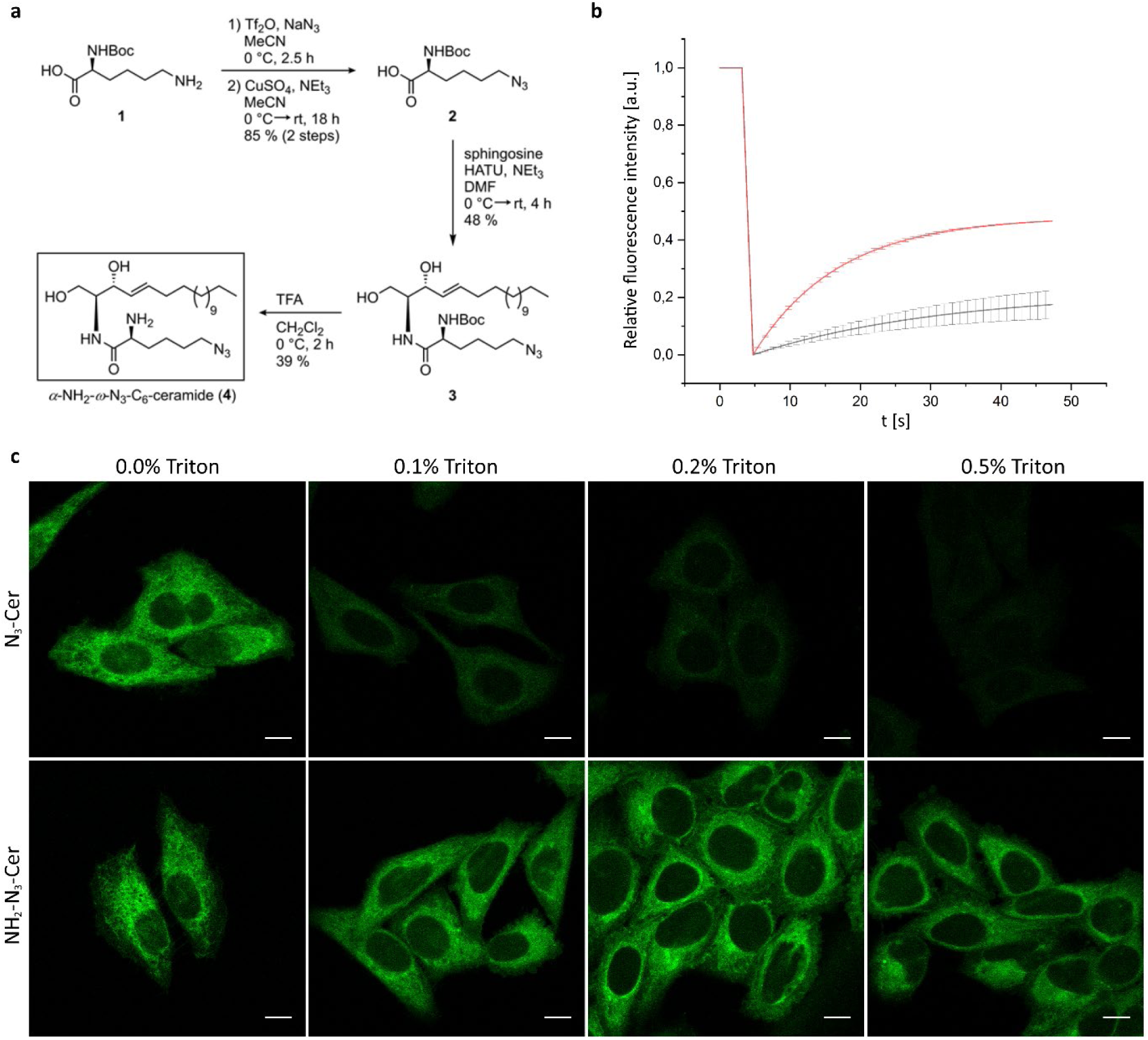
Amino- and azido-functionalized sphingolipids enable fixation and fluorescence labeling of lipids. (a) Schematic overview of the synthesis of α-NH_2_-ω-N_3_-C_6_-Ceramide (for synthesis details see Supporting Information). To investigate the mobility of membrane-incorporated functional sphingolipids HeLa229 cells were fed with 10 µM α-NH_2_-ω-N_3_-C_6_-ceramide or ω-N_3_-C_6_-Ceramide, fixed, permeabilized and stained with DBCO-Alexa Fluor 488. (b) FRAP experiments with the two incorporated ceramide analogues. After three confocal fluorescence imaging frames, a circular region of interest with a diameter 1.8 µm was bleached and fluorescence recovery followed over time. The α-NH_2_-ω-N_3_-C_6_-ceramide (black) shows a lower mobility (mean mobile fraction of 22.2 %) than the ω-N_3_-C_6_-ceramide (red) lacking the primary amino group (mobile fraction of 48.1 %). (c) Confocal fluorescence images of fixed and labeled cells in the presence of increasing concentrations of the detergent Triton-X100. With increasing Triton-X100 concentration ω-N_3_-C_6_-ceramide is efficiently washed out while the α-NH_2_-ω-N_3_-C_6_-ceramide signal remains preserved. Scale bars, 10 µm.

Since glutaraldehyde (GA) can link proteins into hydrogels^9^ we reasoned that α-NH_2_-ω-N_3_-C_6_-ceramides might be as well suited for membrane expansion. To demonstrate its usefulness for ExM we treated HeLa229 with NH_2_-ω-N_3_-C_6_-ceramide followed by glutaraldehyde fixation, permeabilization, fluorescence labeling with DBCO-Alexa Fluor 488, and gelation. For direct comparison we tested the membrane-binding fluorophore-cysteine-lysine-palmitoyl group (mCling), which labels the plasma membrane and is taken up during endocytosis.^23^ Since it carries a primary amine as well, it also remains attached to membranes after fixation and permeabilization and can therefore potentially be used for ExM. In fact, both amino-functionalized membrane probes can be expanded using the GA ExM protocol.^9^ 4x and 10x expanded confocal fluorescence images of ceramide stained cells clearly showed staining of the plasma membrane as well as of membranes of intracellular organelles such as mitochondria, whereas mCling is efficiently incorporated mainly into the cell’s plasma membrane (Figure 2). To verify the expansion factor and investigate if sphingolipid ExM distorts membranes we imaged the same cell before and after 4x and 10x expansion and determined effective expansion factors of 4.1x and 9.8x (Supplementary Figure S15). The confocal fluorescence images of 4x and 10x expanded cellular membranes demonstrate that sphingolipid ExM labeling is dense enough to support nanoscale resolution imaging of continuous membrane structures and even thin membrane protrusions (Figure 2 and Supplementary Figure S15).

**Figure 2.**
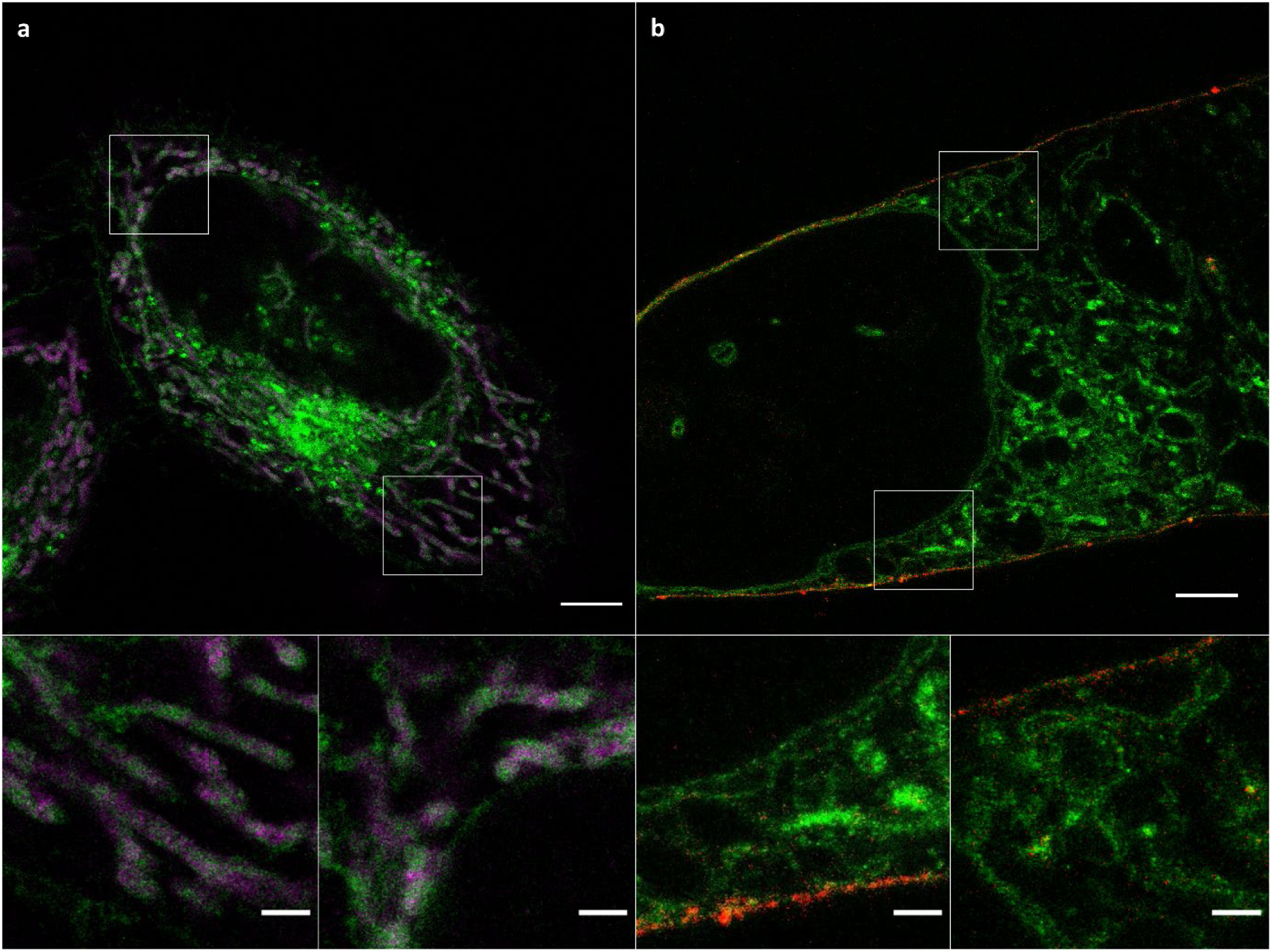
Sphingolipid ExM enables super-resolution imaging of cellular membranes and protein interactions. (a) Confocal fluorescence image of 4x expanded HeLa229 cells. Cells were fed with α-NH_2_-ω-N_3_-C_6_-ceramide, fixed, permeabilized, and labeled with DBCO-Alexa Fluor 488 (green). In addition, Prx3 (magenta), which is located in the mitochondrial matrix was stained by immunolabeling using ATTO 647N labeled secondary antibodies. (b) Confocal fluorescence image of a 10x expanded HeLa229 cell fed with ATTO643-mCling (red) and α-NH_2_-ω-N_3_-C_6_-ceramide clicked with DBCO-Alexa Fluor 488 (green). Scale bars, 20 µm. The images at the bottom show magnified views of the regions outlined by the white boxes in the main images. Scale bars, 5 µm.

### Imaging of expanded lipids and proteins

Furthermore, we tested if the sphingolipid ExM protocol enables imaging of lipids and proteins in the same sample. We therefore immunolabeled the mitochondrial protein Prx3 after permeabilization and click labeling of the bifunctional ceramide. The results obtained clearly showed that the amino-functionalized sphingolipid NH_2_-ω-N_3_-C_6_-ceramide can be used advantageously for super-resolution imaging of cellular membranes and interactions between proteins and ceramides in 4x and 10x expanded samples (Figure 2A). Very recently, Boyden and coworkers introduced an alternative membrane ExM method (mExM) based on a membrane intercalating probe, which enables imaging of 4.5x expanded cellular membranes.^24^ The membrane probe contains a chain of lysines for binding to a polymer anchorable handle and a lipid tail on the amine terminus of the lysine chain, with a glycine in between to provide mechanical flexibility. Furthermore, a biotin residue is attached to enable fluorescence staining of the probe with labeled streptavidin. Both methods, mExM and sphingolipid ExM allow for joint imaging of proteins and lipid membrane structures at nanoscale resolution.

### Sphingolipid ExM of bacterial infections

In addition to the regulation of cellular processes, ceramides play an essential role in infections with pathogenic bacteria.^8,25,26^ These include *Neisseria gonorrhoeae*^*27*^, *Simkania negevensis*^*28*^ and *Chlamydia trachomatis.*^*29,30*^ *C. trachomatis* is by far the best investigated example for an interaction of pathogenic bacterium and host sphingolipid metabolism. This obligate intracellular Gram-negative bacterium is the most frequent cause of bacterial sexually transmitted diseases.^31^ It resides in a membrane-bound vacuole (the inclusion) inside their host cells and undergoes a complex developmental cycle between infectious non-replicating elementary bodies (EB) and non-infectious replicating reticulate bodies (RB). During infection, *Chlamydia* manipulate a plethora of cellular processes, among them the sphingolipid metabolism.^15,16,32^ The ceramide transporter CERT seems to play a key role in ceramide uptake as it strongly localizes in infected cells at the inclusion membrane recruited by the bacterial inclusion protein IncD instead of mediating golgi-ER-trafficking.^33^

To study ceramide uptake by pathogens during infection in more detail we first fed cells with NH_2_-ω-N_3_-C_6_-ceramide for 5 to 60 min 24 h post infection with *C. trachomatis.* The cells were then GA fixed and click-labeled with DBCO-Alexa Fluor 488 for fluorescence imaging. Confocal fluorescence images demonstrated rapid integration of the ceramide into the membrane of *C. trachomatis* already after 5 min and further increasing for longer incubation times (Supplementary Figure S16). This indicates a highly effective and fast ceramide uptake by *C. trachomatis*. Additionally, we applied the specific CERT inhibitor HPA-12 to impede ceramide integration into the bacterial membrane. Fluorescence images recorded after application of HPA-12 showed that HPA-12 efficiently inhibits ceramide uptake by *C. trachomatis* at higher concentrations for short ceramide incubation times of 5 and 15 min (Supplementary Figure S16). For longer ceramide incubation times the influence of HPA-12 treatment on ceramide uptake by bacteria was negligible, suggesting the involvement of different lipid uptake pathways such as vesicle trafficking from the Golgi apparatus.^34^

Next, we investigated if the uptake of ceramides by intracellular pathogens enables ExM of infected cells. Therefore, we fed NH_2_-ω-N_3_-C_6_-ceramide to HeLa229 cells post-infection with *C. trachomatis* and *S. negevensis*, another member of the order *Chlamydiales* (Figure 3). Cells were then fixed with GA, permeabilized, click-labeled with DBCO-Alexa Fluor 488 and expanded using two different ExM protocols. Confocal fluorescence images of the same cells recorded before and after 10x expansion revealed a good quality agreement of bacterial membrane shapes and numbers of bacteria (Supplementary Figure S17). In addition, the post-expansion images clearly showed that the ceramides accumulate strongly in bacterial membranes after infection (Supplementary Figure S17).

**Figure 3.**
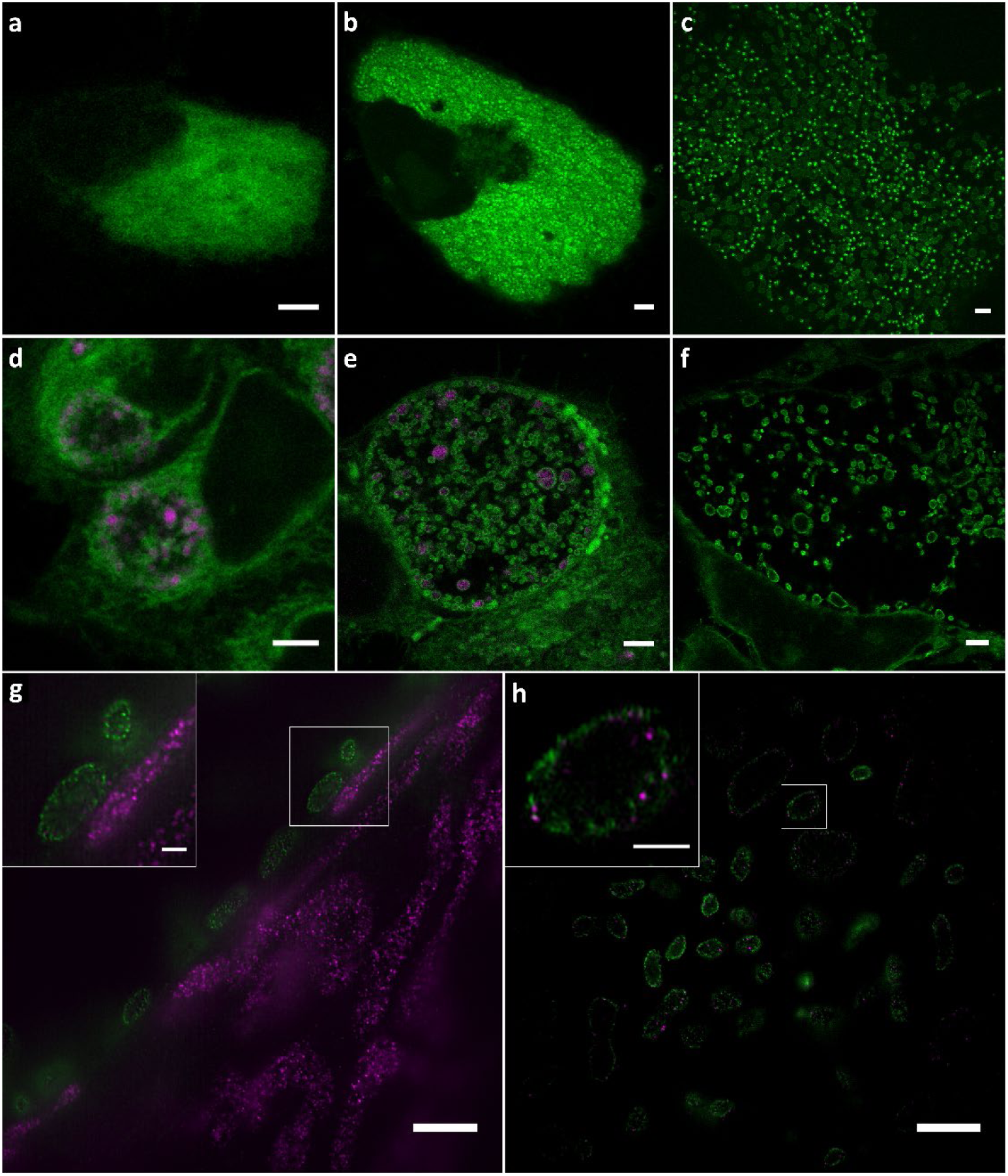
Sphingolipid ExM visualizes intracellular pathogens and their interactions with mitochondrial proteins. (a-c) Cells were infected with *Simkania negevensis* for 96 h, fed with α-NH_2_-ω-N_3_-C_6_-ceramide, fixed, permeabilized and stained with DBCO-Alexa Fluor 488 (green), and then imaged. The images show different cells before expansion (a), after 4x expansion (b), and 10x expansion (c) recorded by confocal microscopy. (d-f) Cells were infected with *Chlamydia trachomatis* for 24 h, fed with α-NH_2_-ω-N_3_-C_6_-ceramide, fixed, permeabilized and stained with DBCO-Alexa Fluor 488 (green). Different cells were imaged before expansion (d), after 4x expansion (e), and 10x expansion (f) by confocal microscopy. In the unexpanded (d) and 4x expanded image (e) chlamydial HSP60 was immulabeled with ATTO647N secondary antibody (magenta). (g) The mitochondrial marker protein Prx3 was stained by immunolabeling with an ATTO 647N secondary antibody (magenta). The confocal fluorescence image of 10x expanded samples revealed a close contact between Chlamydia and mitochondria at the inclusion membrane. (h) SIM images of 10x expanded samples uncover that some Prx3 molecule are inserted into the bacterial membrane. Scale bars, 5 µm (unexpanded images a,d), 10 µm (4x and 10x expanded images b,c,e,f,g,h), and 2 µm (magnified views in images g,h).

Cells infected with a high number of *S. negevensis* required 10x expansion to distinguish individual bacteria (Figure 3A-C). On the other hand, already 4x expansion was sufficient to distinguish between the two forms of *C. trachomatis*, RBs and EBs as has already been shown previously by ExM (Figures 3D,E).^8^ Higher expansion (10x ExM) demonstrated that the ceramide signal accumulates in the membranes of the two pathogens *C. trachomatis* and *S. negevensis* (Figure 3C,F). The fluorescence signals of host cell membranes appeared comparably dim (compare Figure 2 and Figure 3) indicating an extremely efficient ceramide uptake by bacteria. Corresponding control experiments with ω-N_3_-C_6_-ceramide and DBCO-Alexa Fluor 488 alone showed only very weak background staining (Supplementary Figure S18). These results clearly show that sphingolipid ExM enables very dense and continuous membrane staining of intracellular bacteria and thus imaging of bacterial membranes with a resolution hitherto only provided by electron microscopy.

So far, we focused our investigations on bacterial infections and demonstrated the fast and efficient incorporation of NH_2_-ω-N_3_-C_6_-ceramide into *C. trachomatis* and *S. negevensis* and their 4x and 10x expansion (Figure 3). The introduced method can be highly valuable for studying not only host pathogen interactions but also lipid metabolism. It would be very interesting to investigate the interaction of Inc proteins involved in ceramide transport like IncD, CERT, as well as Golgi and ER proteins to further elucidate the uptake and incorporation of ceramides into the membrane of *C. trachomatis*.

Furthermore, we tested if the ceramide underlying structure sphingosine can be used successfully for ExM. The sphingoid base backbone sphingosine carries a natural amino group and plays a central role in infections with *N. gonorrhoeae* among other bacterial pathogens.^35^ Addition of ω-N_3_-sphingosine to infected Chang cells followed by GA fixation, permeabilization, click labelling with DIBO-Alexa Fluor 488 and gelation demonstrated the general applicability of the method. Details of intracellular *N. gonorrhoeae* can be clearly visualized by sphingolipid ExM (Supplementary Figure S19).

### Imaging interactions of bacteria and intracellular proteins

To demonstrate the compatibility of sphingolipid ExM for investigations of pathogen interactions with intracellular proteins, we investigated chlamydial interactions with mitochondria. It is known that *C. trachomatis* reorganizes the host organelles. However, so far all investigations have been performed by confocal fluorescence imaging or electron microscopy.^36^ Hence, we immunolabeled the mitochondrial matrix protein Prx3 and incorporated ceramides in *C. trachomatis* infected cells before gelation. The corresponding confocal fluorescence images of 10x expanded samples showed the mitochondrial rearrangement after infection with *C. trachomatis* as mitochondria localized around the inclusion (Figure 3G). To highlight details of this interaction by a higher spatial resolution we used structured illumination microscopy (SIM)^37^, which allowed us to uncover direct interactions between mitochondria and *C. trachomatis* (Figure 3H). In some cases, Prx3 signals appeared to be located in bacteria indicating unspecific protein uptake. Similar experiments performed in the absence of primary antibodies demonstrated that the signals detected in bacteria are not caused by nonspecific binding of the used secondary antibody (Supplementary Figure S20). Albeit ceramides accumulate strongly in bacterial membranes the labeling density of intracellular membranes is still high enough to enable nanoscale imaging of protein-pathogen interactions in infected cells.

Interestingly, we could often detect individual Chlamydia within close proximity to the inclusion membrane after feeding with NH_2_-ω-N_3_-C_6_-ceramides, possibly indicating an active docking to the inclusion membrane and an absorption of nutrition by *C. trachomatis* (Supplementary Figure S21 and Supplementary Movie 1) as has been hypothesized earlier^38^ and reported in electron microscopy studies.^36,39^ This behavior has previously been proposed as a mechanism by which RBs acquire nutrients including host lipids^40^ and as an essential step in chlamydial development.^38^ However, previous attempts to localize chlamydial particles in the inclusion required highly laborious techniques such as Serial block-face scanning electron microscopy.^36^ Using sphingolipid ExM with clickable probes, the three-dimensional structure of lipid interfaces can be imaged at a lateral resolution of ∼20 nm by confocal fluorescence microscopy. This enables to investigate metabolite acquisition like highly efficient ceramide transfer from the host to bacterial membranes at higher resolution and even in three dimensions.

### 10x Sphingolipid ExM-SIM resolves the double membrane of intracellular bacteria

Whereas transport of ceramide to the *Chlamydia* inclusion has been reported earlier^29^, one of the unanswered questions is whether ceramides form parts of the bacterial outer (OM) or inner membrane (IM) or of both these membranes. Indeed, SIM images of 10x expanded chlamydia demonstrated that NH_2_-ω-N_3_-C_6_-ceramides are efficiently incorporated into the IM and OM of intracellular *Chlamydia* (Figures 4A,B). The high labeling efficiency in combination with the high spatial resolution of 10-20 nm of 10x expanded samples provided by SIM allowed us to resolve the IM and OM (Figure 4). We investigated three different infected cells and selected those bacteria whose orientation allowed us to visualize spatially separated OM and IM (i.e. frontal views of bacteria) and determine the distance between the two membranes to 27.6 ± 7.7 nm (s.d.) from 23 cross sectional intensity profiles (Supplementary Figures S22 and S23). This value is typical for the separation of OM and IM of gram-negative bacteria and in agreement with electron microscopy data.^41^ Since the mechanism of bacterial membrane biogenesis from host-derived lipids is currently unknown, our findings of the ceramide incorporation in both bacterial membranes suggest an active process rather than only the fusion of lipid vesicles with the surface and exclusive integration into the outer membrane of *Chlamydia*.

**Figure 4.**
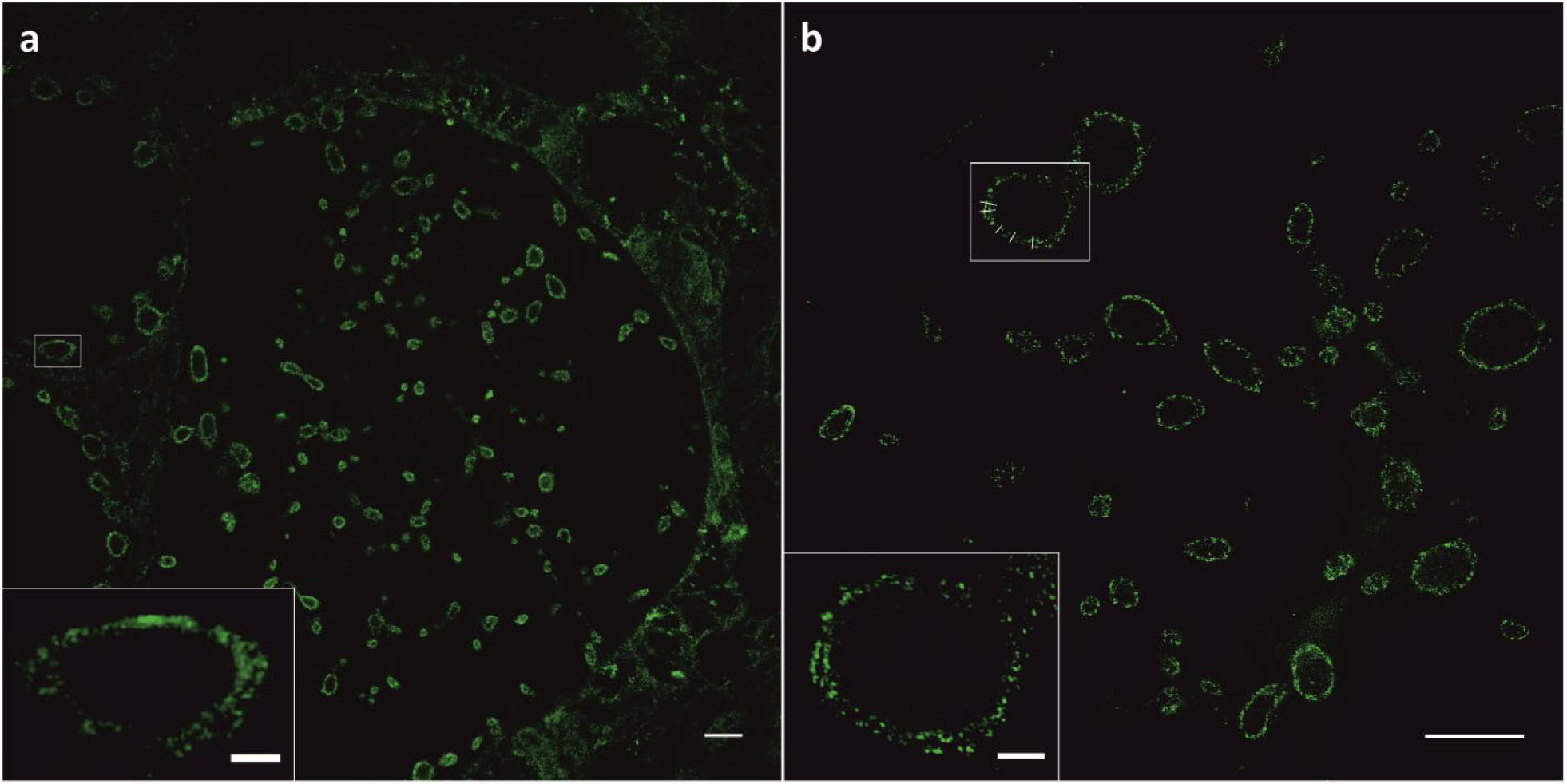
10x Sphingolipid ExM in combination with SIM resolves the distance between the OM and IM of gram-negative bacteria. HeLa229 cells infected with *Chlamydia trachomatis* for 24 h, fed with α-NH_2_-ω-N_3_-C_6_-ceramide, fixed, permeabilized and click-labeled with DBCO-Alexa Fluor 488 (green). Confocal (a) and SIM images (b) disclose that ceramides are incorporated into the OM and IM. Fitting intensity cross sectional profiles at different positions by a bimodal Gaussian fit resulted in a peak-to-peak distance of 27.6 ± 7.7 nm (s.d.) (Supplementary Figures S22 and S23). Scale bars 10 µm (a,b), 2 µm (white boxes).

The high spatial resolution provided by sphingolipid ExM may also be used to study mechanisms of antibiotic resistance. Infections with multidrug-resistant gram-negative bacteria are difficult to treat because of the double membrane that is impermeable for most antibiotics.^42^ Hence, being able to visualize the double membrane might promote the development of antibiotics with improved membrane permeability. Furthermore, sphingolipid ExM can also be used advantageously to investigate various ceramide pathways related to apoptosis, proliferation, cancer, inflammation, and neurodegeneration.^34,43^

## Conclusions

ExM has facilitated super-resolution imaging of cells and tissues with standard fluorescence microscopes available in most research facilities, yet it has been limited to the expansion of proteins and nucleic acids due to the lack of primary amino groups in lipids. We have developed the double-functionalized sphingolipid NH_2_-ω-N_3_-C_6_-ceramide that incorporates efficiently into cellular and bacterial membranes and can be fixed, fluorescently labeled by click chemistry, and linked into polyelectrolyte hydrogels by GA treatment. The mechanism by which GA fixes and crosslinks amino-modified ceramides into hydrogels is less obvious but most probably associated with the existence of multimeric forms of GA containing aldehyde and alkene groups, which both can potentially be covalently linked to the acrylamide polymer.^9^ Sphingolipid ExM allows for simultaneous super-resolution imaging of membranes and associated proteins in 4x and 10x expanded samples. In combination with SIM, sphingolipid ExM enables 10-20 nm spatial resolution, approaching that of electron microscopy and has allowed us to resolve details of sphingolipid-protein interactions. Such high spatial resolutions are difficult to achieve using pre-expansion immunolabeling with primary and secondary antibodies but feasible using small membrane incorporated ceramides that are linked into the polymer and fluorescently labeled with minimal linkage error. For clarification, pre-expansion immunolabeling introduces a linkage error of ∼ 17.5 nm^7^, which translates into a linkage error of ∼ 175 nm after 10x expansion. Such large linkage errors severely blur the underlying structure and impede super-resolution imaging with a spatial resolution of 10-20 nm in 10x expanded samples. We hypothesize that our approach of introducing a primary amino group for fixation and linkage into acrylamide polymers by GA can be broadly used to enable ExM of other lipids and thus far inaccessible molecule classes including carbohydrates.

## Methods

### Chemical Synthesis of *α*-Amino-*ω*-Azido-C_6_-Ceramide

Starting from *N*-Boc-protected L-lysine (**1**) the introduction of the azide-functionality was accomplished *via* catalytic diazotransfer reaction to obtain azido-acid **2** in 85 % yield. For that, triflyl azide was prepared based on a method of *Yan et al*. with a reduced amount of highly toxic sodium azide and triflyl anhydride compared to previous protocols.^44^ Subsequent amide coupling of **2** with sphingosine was performed in DMF under basic conditions using HATU as coupling reagent. The resulting Boc-protected azido-ceramide analogue **3** was isolated in 48 % yield. In the last step the amine group was deprotected by the treatment with TFA in dichloromethane. After basic workup, followed by column chromatography, the target ceramide analogue **4** was successfully isolated in 39% yield. Details on the experimental procedures can be found in the Supporting Information. All isolated compounds were characterized by a combination of HRMS, NMR and IR spectroscopy (Supplementary Figures S1-S13).

### Cell lines and bacteria

Human HeLa229 cells (ATCC CCL-2.1tm) and human epithelial conjunctival cells (Chang) were cultured in 10 % (v/v) heat inactivated FBS (Sigma-Aldrich) RPMI1640 + GlutaMAXtm medium (Gibcotm) and were grown in a humidified atmosphere containing 5 % (v/v) CO_2_ at 37 °C. HeLa229 cells were used for infection with *Chlamydia trachomatis* and *Simkania negevensis*, Chang cells for infection with *Neisseria gonorrhoeae.* For this study, *C. trachomatis* serovar L2/434/Bu (ATCC VR-902B™), *S. negevensis* and *N. gonorrhoeae* (strain MS11, derivative N927) were used. *C. trachomatis* and *S. negevensis* were cultivated as previously described.^8,28^ For this, the bacteria were propagated in HeLa229 cells at a multiplicity of infection (MOI) of 1 for 48 h for *C. trachomatis* and 72 h for *S. negevensis*. The cells were then detached and lysed using glass beads (3 mm, Roth). Low centrifugation supernatant (10 min at 2000 g at 4 °C for *C. trachomatis* and 10 minutes at 600 g at 4 °C for *S. negevensis*) was transferred to high speed centrifugation (30 min at 30.000 g at 4 °C for *C. trachomatis* and 30 min at 20.000 g at 4 °C for *S. negevensis*) to pellet the bacteria. Afterwards, the pellet was washed and resuspended in 1x SPG buffer (7.5 % sucrose, 0.052 % KH_2_PO_4_, 0.122 % NaHPO_4_, 0.072 % L-glutamate). The resuspended bacteria were then stored at -80 °C and titrated for an MOI of 1 for further experimentation. Infected cells were incubated in a humidified atmosphere with 5 % (v/v) CO_2_ at 35 °C. The cell lines as well as the Chlamydia used in this study were tested to be free of Mycoplasma *via* PCR. Neisseria were cultivated on gonococci (GC) agar (ThermoScientific, Waltham, USA) plates supplemented with 1 % vitamin mix at 37 °C and 5 % CO_2_ for 16 h. On the day of infection, liquid culture was performed in protease-peptone medium (PPM) supplemented with 1 % vitamin mix and 0.5 % sodium bicarbonate 8.4 % solution (PPM+) at 37 °C and 120 rpm. Gonococci were grown to an OD_550_ 0.4 to 0.6. Before infecting the cells, the medium of the liquid culture was changed to 4-(2-Hydroxyethyl)piperazine-1-ethanesulfonic acid (HEPES buffer) medium by centrifugation with 4000 rpm for 5 min. After the indicated time of 4 h, the infection was stopped by washing the cells three times with Hepes medium.

### Click-Chemistry and Immunolabeling

For immunostaining, cells were seeded on 15 mm coverslips. α-Amino-ω-Azido-C_6_-Ceramide, ω-Azido-C_6_-Ceramide, as well as ω-Azido-Sphingosine were fed with 10 µM final concentration for 1 h at 37 °C. For chlamydial infection, the cells were fed with ceramide-analogues 23 h post infection and for infection with *Simkania for 72 h and* for neisserial infection the cells were fed with the sphingosine analogue immediately before infection. Afterwards, the cells were fixed in 4 % PFA and 0.1 % GA for 15 min, washed 3x in 1xPBS and then permeabilized for 15 min in 0.2 % Triton X-100 in PBS. The cells were then washed again 3x in 1xPBS and then incubated with 5 µM DBCO-488 (Jena Bioscience, CLK-1278-1) at 37 °C for 30 min or 5 µM Click-IT Alexa Fluor® 488 DIBO alkyne dye (ThermoScientific, Waltham, USA) at 37 °C for 30 min. For staining with antibodies, the cells were washed, blocked using 2 % FCS in 1xPBS for 1 h and then incubated in primary antibody diluted in blocking buffer for 1 h in a humid chamber. The primary antibodies used in this study were: anti-HSP60 ms (Santa Cruz, sc-57840, dilution 1:200), anti-*Neisseria gonorrhoeae* primary antibody rb (US biological, dilution 1:200), anti-Prx3 (Origene, TA322470, dilution 1:100) and anti-CERT (Abcam, ab72536, 1:100). After that, the cells were washed 3x in 1xPBS and then incubated in the corresponding secondary antibody diluted in blocking buffer for 1 h and then washed 3x with 1xPBS. The secondary antibodies used were: ATTO 647N ms (Rockland, 610-156-121S, dilution 1:200) and ATTO 647N rb (Sigma, 40839, dilution 1:200).

### mCling

mCling (Biosyntan) was labeled as previously published.^23^ In short, 150 nmol mCling was incubated in 3 molar excess of ATTO 643-Maleimide (ATTO-TEC, AD 643-45) in 100 mM TCEP overnight at RT under continuous shaking. The label product was purified by HPLC (JASCO) and the concentration was determined using a UV-vis spectrophotometer (Jasco V-650). Staining with mCling was performed by the incubation of living cells in 0.5 µM mCling dissolved in media for 10 min at 37°C.

### Expansion Microscopy

Stained cells were treated according to Kunz *et al.*^8^ for 10 min with 0.25 % GA at RT and gelated after three washing steps. In case of 4x expansion a monomer solution consisting of 8.625 % sodium acrylate (Sigma, 408220), 2.5 % acrylamide (Sigma, A9926), 0.15 % N,N’-methylenbisacrylamide (Sigma, A9926), 2 M NaCl (Sigma, S5886) and 1xPBS and 0.2 % freshly added ammonium persulfate (APS, Sigma, A3678) and tetramethylethylenediamine (TEMED, Sigma, T7024) was used. Here gelation was performed for 1 h at RT followed by proteinase digestion. In case of 10x expansion 1 ml of the monomer solution containing 0.267 g DMAA (Sigma, 274135) and 0.064 g sodium acrylate (Sigma, 408220) dissolved in 0.57 g ddH_2_O was degassed for 45 min on ice with nitrogen followed by the addition of 100 µl KPS (0.036 g/l, Sigma, 379824). After another 15 min of degassing and the addition of 4 µl TEMED per ml monomer solution, gelation was performed for 30 min at RT followed by an incubation of 1.5 h at 37 °C. Hereafter the samples were digested for 3 h – overnight in digestion buffer (50 mM Tris pH 8.0, 1 mM EDTA (Sigma, ED2P), 0.5 % Triton X-100 (Thermo Fisher, 28314) and 0.8 M guanidine HCl (Sigma, 50933)), supplied with 8 U/ml protease K (Thermo Fisher, AM2548) and for expansion of Neisseria additional 1 mg/ml Lysozyme according to Lim *et al.*^45^ Digested gels were expanded in hourly changed ddH_2_O until the expansion saturated. The expansion factor was determined by the gel size using calipers directly after gelation and by the gel size of the digested and expanded samples. We achieved experimental expansion factors of 4.1 for the 4x monomer solution and 10 for the 10x monomer solution, and the expansion factor remained constant for the used monomer solutions. Expanded and chopped gels were stored at 4°C in ddH_2_O immobilized prior to imaging on PDL-coated glass chambers (Merck, 734-2055).

### Confocal Microscopy and SIM

Confocal imaging was performed on an inverted microscope (Zeiss LSM700) or on a Leica TCS SP5 confocal microscope (Leica Biosystems) and SIM-imaging on a Zeiss ELYRA S.1 SR-SIM structured illumination platform using a 63x water-immersion objective (C-Apochromat, 63x 1.2 NA, Zeiss, 441777-9970). Reconstruction of SIM-images was performed using the ZEN image-processing platform with a SIM module. Z-stacks were processed using Imaris 8.4.1 and FIJI 1.51n.^46^

### FRAP

HeLa229 cells were seeded in an 8-well chambered high precision coverglass (Sarstedt 8-well on coverglass II) and incubated for 24 h at 37 °C and 5 % CO_2_. The cells were fed with 10 µM of the corresponding azido-ceramide analogue for 30 min in cell culture media. Afterwards, the cells were washed with HBSS with magnesium and calcium and fixed with 4 % formaldehyde and 0.1 % glutaraldehyde in HBSS for 15 min at room temperature and washed. Ceramides were labelled by strain-promoted alkyne-azide cycloaddition (SPAAC) with 10 µM DBCO-Alexa Fluor 488 in HBSS for 30 min at 37 °C and washed. FRAP-imaging was performed at a confocal laser scanning microscope (CLSM) LSM700 (Zeiss, Germany) using the Plan-Apochromat 63x 1.4 oil objective. Using the 488 nm laser line as excitation, a time series with 30 frames every 1.5 s was recorded. After three frames a circular region of interest with diameter 1.8 µm was bleached and fluorescence recovery followed over time.

## Associated content

The Supporting Information is available free of charge on the ACS Publications website.

Supplementary Figures S1-S23.

Supplementary Movie 1: 10x ExM SIM z-stack of Hela229 cells infected with *Chlamydia trachomatis* for 24 h, fed with α-NH_2_-ω-N_3_-C_6_-ceramide, fixed, permeabilized and stained with DBCO-Alexa Fluor 488. Chlamydia are clearly located at the inclusion membrane. Scale bar, 10 µm.

## Author Contributions

The manuscript was written through contributions of all authors. R.G. and T.C.K. designed and performed experiments, analyzed data and wrote the manuscript. J.F. and J.S. synthesized α-Amino-ω-Azido-C6-Ceramide, J.Sch., F.S. and V.K-P. performed experiments. T.R. and M.S. designed the experiments and wrote the manuscript. All authors have given approval to the final version of the manuscript. ‡These authors contributed equally.

## Acknowledgments

We thank Elke Maier for the preparation of *S. negevensis* stocks. This work was supported by the Deutsche Forschungsgemeinschaft (DFG) GRK 2157 to VKP, TR and MS, and DFG FOR 2123 to TR and MS.

## Supporting Information

### Synthetic Procedure and Characterization

#### *N*^2^-(*tert*-butoxycarbonyl)-*N*^6^-diazo-L-lysine (2)

To a suspension of NaN_3_ (380 mg, 5.85 mml, 1.44 eq.) in MeCN (5 mL) was added Tf_2_O (820 μL, 4.87 mmol, 1.20 eq.) dropwise at 0 °C. The mixture was stirred for 2.5 h at 0 °C and the resulting solution was then transferred to a mixture of (*tert*-butoxycarbonyl)-L-lysine (**1**) (1.00 g, 4.06 mmol, 1.00 eq.), NEt_3_ (1.13 mL, 8.12 mmol, 2.00 eq.) and CuSO_4_ (6.48 mg, 40.6 μmol, 0.01 eq.) in MeCN (10 mL) at 0 °C. The ice bath was removed and the reaction mixture was stirred at RT for 18 h. After the addition of H_2_O (50 mL) and 1 M aq. HCl (20 mL), the aqueous phase was extracted with EtOAc (4 x 60 mL). The combined organic phases were dried (MgSO_4_) and concentrated under reduced pressure. The yellow residue was purified by column chromatography on silica gel (CH_2_Cl_2_/MeOH 20:1) to give **2** (936 mg, 3.44 mmol, 85 %) as a colourless oil, which partly crystallized.^1^

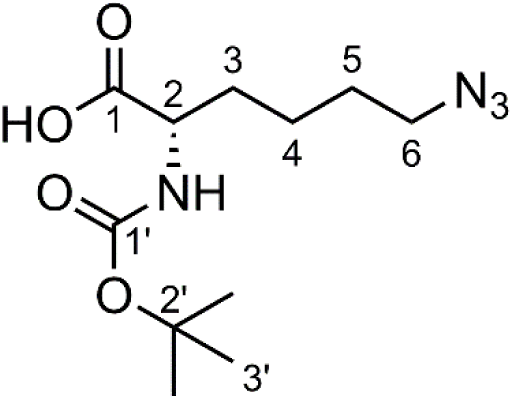

**Formula**: C_11_H_20_N_4_O_4_ (272.31 g/mol).

***R***_**f**_ (CH_2_Cl_2_/MeOH 30:1): 0.27.

^**1**^**H NMR** (CDCl_3_, 400 MHz): *δ* (2 rotamers) = 1.45 (s, 9H, *H*-3’), 1.46–1.57 (m, 2H, *H*-4), 1.57–1.66 (m, 2H, *H*-5), 1.66–1.80 (m, 1H, *H*-3), 1.80–1.96 (m, 1H, *H*-3), 3.29 (t, ^3^*J*_6,5_ = 6.7 Hz, 2H, *H*-6), 4.07–4.40 (m, 1H, *H*-2), 5.06/6.55 (each br d, ^3^*J*_NH,2_ = 8.1/5.5 Hz, together 1H, N*H*), 9.93 (br s, 1H, O*H*) ppm.

^**13**^**C NMR** (CDCl_3_, 100 MHz): *δ* (2 rotamers) = 22.7 (*C*-4), 28.4 (3C, *C*-3’), 28.5 (*C*-5), 32.0/32.2 (*C*-3), 51.2 (*C*-6), 53.3/54.5 (*C*-2), 80.6/82.1 (*C*-2’), 155.8/157.0 (*C*-1’), 176.8/177.4 (*C*-1) ppm.

**HRMS** (ESI^+^): *m/z* calcd. for C_11_H_20_N_4_NaO_4_ [M+Na]^+^: 295.1377; found: 295.1383 (|Δ*m/z*| = 2.0 ppm); *m/z* calcd. for C_22_H_40_N_8_NaO_8_ [2M+Na]^+^: 567.2861; found: 567.2866 (|Δ*m/z*| = 0.8 ppm).

**FTIR** (ATR): 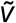 = 3313, 3186, 3073, 2979, 2936, 2870, 2095, 1712, 1686, 1514, 1479, 1456, 1392, 1368, 1281, 1234, 1189, 1150, 1106, 1051, 1024, 940, 852, 779, 753, 736, 643, 608, 564 cm^−1^.

The measured spectroscopic data are in agreement with previously reported data.^2-5^

#### *tert*-Butyl ((*S*)-6-azido-1-(((2*S*,3*R,E*)-1,3-dihydroxyoctadec-4-en-2-yl)amino)-1-oxohexan-2-yl)carbamate (3)

To a solution of azido-acid **2** (45.5 mg, 167 μmol, 1.00 eq.) in dry DMF (3 mL) were added NEt_3_ (69.9 μL, 501 μmol, 3.00 eq.) and HATU (69.8 mg, 184 μmol, 1.10 eq.) at 0 °C. After stirring at this temperature for 20 min, sphingosine (50.0 mg, 167 μmol, 1.00 eq.) and dry DMF (3 mL) were added. The ice bath was removed and the reaction mixture was stirred at rt for 3.5 h. After the addition of H_2_O (10 mL) and saturated aq. NH_4_Cl solution (30 mL), the aqueous layer was extracted with EtOAc (5 x 20 mL). The combined organic phases were washed with brine (20 mL), dried (MgSO_4_) and concentrated under reduced pressure. The oily residue was purified by column chromatography on silica gel (CHCl_3_/MeOH 40:1 to 30:1) to give **3** (44.1 mg, 79.6 μmol, 48 %) as a colourless, waxy solid.

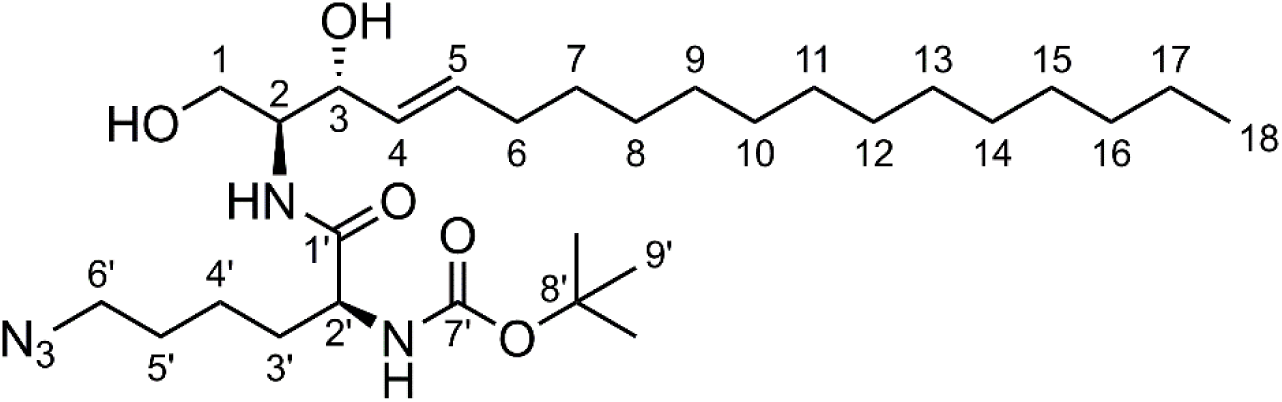

**Formula**: C_29_H_55_N_5_O_5_ (553.79 g/mol).

***R***_**f**_ (CH_2_Cl_2_/MeOH 30:1): 0.15.

^**1**^**H NMR** (CDCl_3_, 400 MHz): *δ* = 0.88 (t, ^3^*J*_18,17_ = 6.9 Hz, 3H, *H*-18), 1.25–1.31 (m, 20H, *H*-8–17), 1.35–1.38 (m, 2H, *H*-7), 1.44 (s, 9H, *H*-9’), 1.48–1.90 (m, 6H, *H*-3’–5’), 2.03–2.08 (m, 2H, *H*-6), 3.29 (t, ^3^*J*_6’,5’_ = 6.7 Hz, 2H, *H*-6’), 3.71 (dd, ^2^*J*_1,1_ = 11.6 Hz, ^3^*J*_1,2_ = 3.3 Hz, 1H, *H*-1), 3.83–3.86 (m, 1H, *H*-2), 3.96 (dd, ^2^*J*_1,1_ = 11.6 Hz, ^3^*J*_1,2_ = 3.3 Hz, 1H, *H*-1), 3.99– 4.04 (m, 1H, *H*-2’), 4.36–4.38 (m, 1H, *H*-3), 5.07 (d, ^3^*J*_NH,2’_ = 6.9 Hz, 1H, N*H*), 5.51 (ddt, ^3^*J*_4,4_ = 15.4 Hz, ^3^*J*_4,3_ = 6.0 Hz, ^4^*J*_4,6_ = 1.3 Hz, 1H, *H*-4), 5.80 (dtd, ^3^*J*_5,4_ = 15.4 Hz, ^3^*J*_5,6_ = 6.8 Hz, ^4^*J*_5,3_ = 1.4 Hz, 1H, *H*-5), 6.86 (d, ^3^*J*_NH,2_ = 8.0 Hz, 1H, N*H*) ppm.

^**13**^**C NMR** (CDCl_3_, 100 MHz): *δ* = 14.3 (*C*-18), 22.8 (*C*-17), 23.0 (*C*-4’), 28.4 (3C, *C*-9’), 28.6 (*C*-5’), 29.3 (*C*-7), 29.4, 29.5, 29.6, 29.8, 29.8, 29.8 (8C, *C*-8–15), 32.0 (2C, C-3’ & C-16), 32.5 (*C*-6), 51.3 (*C*-6’), 54.6 (*C*-2), 55.1 (*C*-2’), 61.9 (*C*-1), 74.0 (*C*-3), 80.7 (*C*-8’), 128.6 (*C*-4), 134.2 (*C*-5), 156.2 (*C*-7’), 172.5 (*C*-1’) ppm.

**HRMS** (ESI^+^): *m/z* calcd. for C_29_H_55_N_5_NaO_5_ [M+Na]^+^: 576.4095; found: 576.4073 (|Δ*m/z*| = 3.8 ppm); *m/z* calcd. for C_58_H_110_N_10_NaO_10_ [2M+Na]^+^: 1129.8299; found: 1129.8274 (|Δ*m/z*| = 2.2 ppm).

#### (*S*)-2-Amino-6-azido-*N*-((2*S*,3*R,E*)-1,3-dihydroxyoctadec-4-en-2-yl)hexanamide / *α*-NH_2_-*ω*-N_3_-C_6_-ceramide (4)

To a solution of carbamate **3** (37.0 mg, 6.68 μL) in CH_2_Cl_2_ (1 mL) was added TFA (200 μL) at 0 °C. The reaction mixture was stirred at this temperature for 2 h and was then quenched by the addition of H_2_O (5 mL) and 1 M aq. NaOH (8 mL). After the extraction with EtOAc (5 x 10 mL), the combined organic phases were washed with brine (10 mL), dried (MgSO_4_) and the solvent was removed under reduced pressure. The residue was purified by column chromatography on silica gel (CHCl_3_/MeOH/25 % aq. NH_3_ 9:1:0.1) to give the crude product containing some NH_3_ salts. To remove these impurities, the residue was dissolved in CH_2_Cl_2_ (15 mL) and washed with saturated aq. NaHCO_3_ solution (10 mL). The aqueous phase was extracted with CH_2_Cl_2_ (6 x 5 mL), washed with brine (5 mL) and dried (MgSO_4_). The solvent was removed under reduced pressure to give **4** (11.9 mg, 26.2 μmol, 39 %) as a colourless solid.

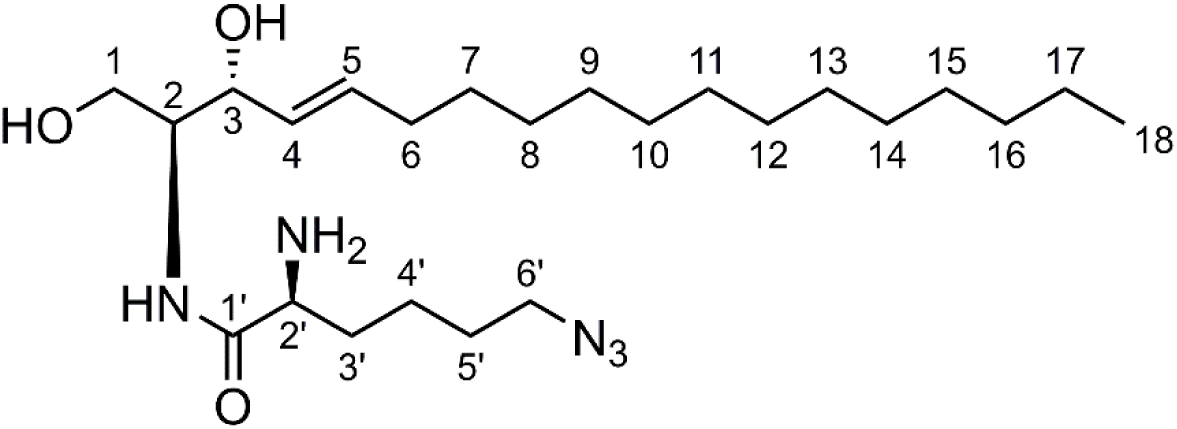

**Formula**: C_24_H_47_N_5_O_3_ (453.67 g/mol).

***R***_**f**_ (CHCl_3_/MeOH/25 % aq. NH_3_ 9:1:0.1):0.26.

^**1**^**H NMR** (CDCl_3_, 600 MHz): *δ* = 0.87 (t, ^3^*J*_18,17_ = 7.1 Hz), 1.25–1.30 (m, 20H, *H*-8–17), 1.33–1.37 (m, 2H, *H*-7), 1.43–1.52 (m, 2H, *H*-4’), 1.53–1.59 (m, 1H, *H*-3’), 1.59–1.67 (m, 2H, *H*-5’), 1.83–1.88 (m, 1H, *H*-3’), 2.03–2.07 (m, 2H, *H*-6), 2.21 (br s, 4H, 2 x O*H* & N*H*_2_), 3.26–3.33 (m, 2H, *H*-6’), 3.39 (dd, ^3^*J*_2’,3’_ = 7.9 Hz, ^3^*J*_2’,3’_ = 4.7 Hz, 1H, *H*-2’), 3.71 (dd, ^2^*J*_1,1_ = 11.4 Hz, ^3^*J*_1,2_ = 3.5 Hz, 1H, *H*-1), 3.83–3.86 (m, 1H, *H*-2), 3.91 (dd, ^2^*J*_1,1_ =11.4 Hz, ^3^*J*_1,2_ =4.3 Hz, 1H, *H*-1), 4.30–4.32 (m, 1H, *H*-3), 7.82 (d, ^3^*J*_NH,2_ = 7.7 Hz, 1H, N*H*), 5.52 (ddt, ^3^*J*_4,4_ = 15.4 Hz, ^3^*J*_4,3_ = 6.5 Hz, ^4^*J*_4,6_ = 1.4 Hz, 1H, *H*-4), 5.78 (dtd, ^3^*J*_5,4_ = 15.4 Hz, ^3^*J*_5,6_ = 6.8 Hz, ^4^*J*_5,3_ = 1.2 Hz, 1H, *H*-5) ppm.

^**13**^**C NMR** (CDCl_3_, 150 MHz): *δ* = 14.3 (*C*-18), 22.8 (*C*-17), 23.1 (*C*-4’), 28.8 (*C*-5’), 29.3 (*C*-7), 29.4, 29.5, 29.6, 29.8, 29.8, 29.8, 29.8 (8C, *C*-8–15), 32.1 (*C*-16), 32.5 (*C*-6), 34.8 (*C*-3’), 51.3 (*C*-6’), 55.0 (*C*-2), 55.3 (*C*-2’), 62.7 (*C*-1), 74.3 (*C*-3), 128.8 (*C*-4), 134.5 (*C*-5), 175.7 (*C*-1’) ppm.

^**15**^**N NMR** (CDCl_3_, 60 MHz): *δ* = −352.3 (*N*H_2_), −309.4 (C-*N*-N-N), −265.8 (*N*H), −132.8

(C-N-*N*-N) ppm.

**HRMS** (ESI^+^): *m/z* calcd. for C_24_H_47_N_5_NaO_3_ [M+Na]^+^: 476.35711; found: 476.35763 (|Δ*m/z*| = 1.08 ppm).

**FTIR** (ATR): 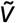 = 3414, 3315, 3271, 3183, 3087, 2677, 2096, 1651, 1614, 1552, 1467, 1455, 1438, 1365, 1348, 1304, 1249, 1165, 1126, 1082, 1044, 1012, 992, 962, 932, 822, 801, 720, 801, 720, 650, 616, 590, 557 cm^−1^.

## NMR Spectra

**Supplementary Figure S1.**
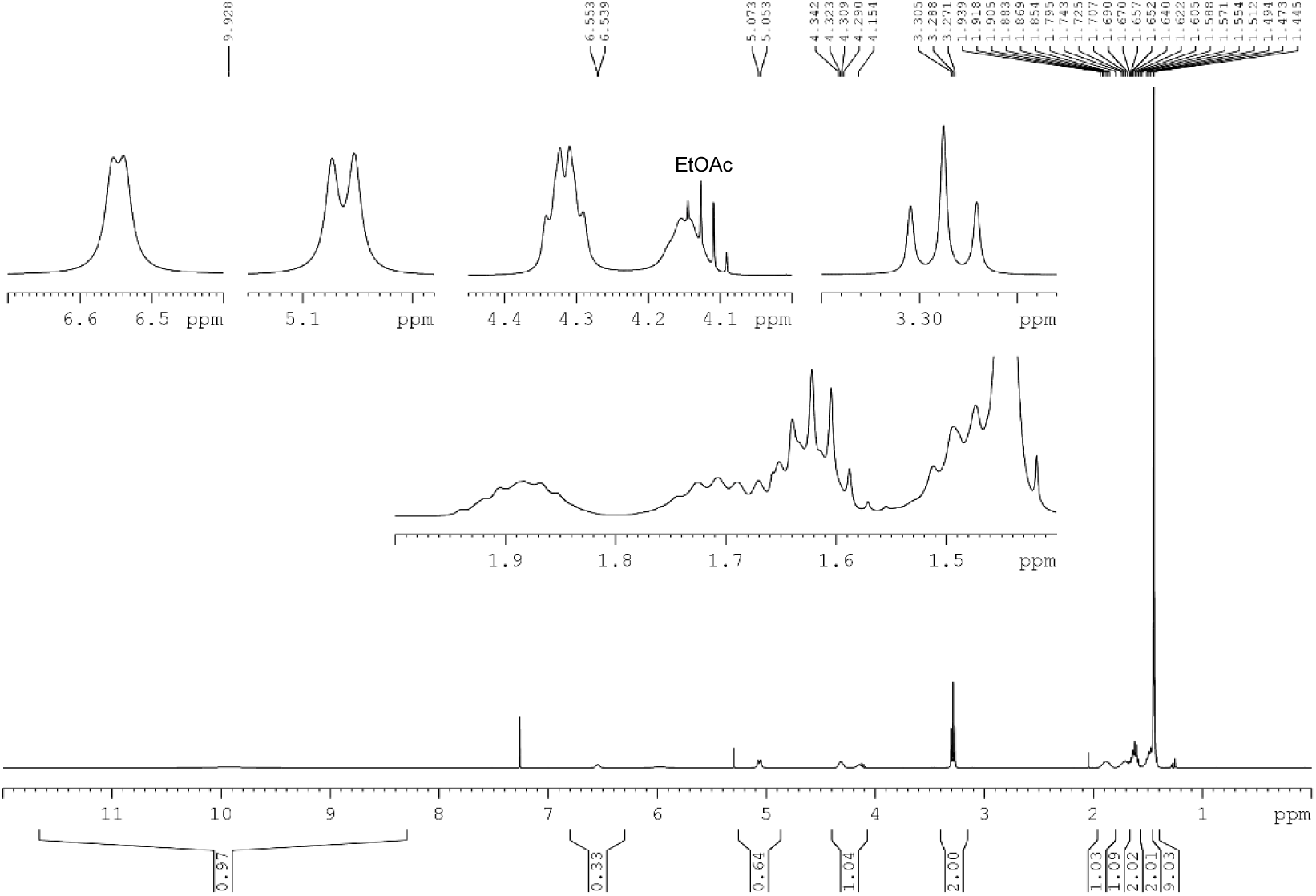
^1^H NMR spectrum of **2** (CDCl_3_, 400 MHz).

**Supplementary Figure S2.**
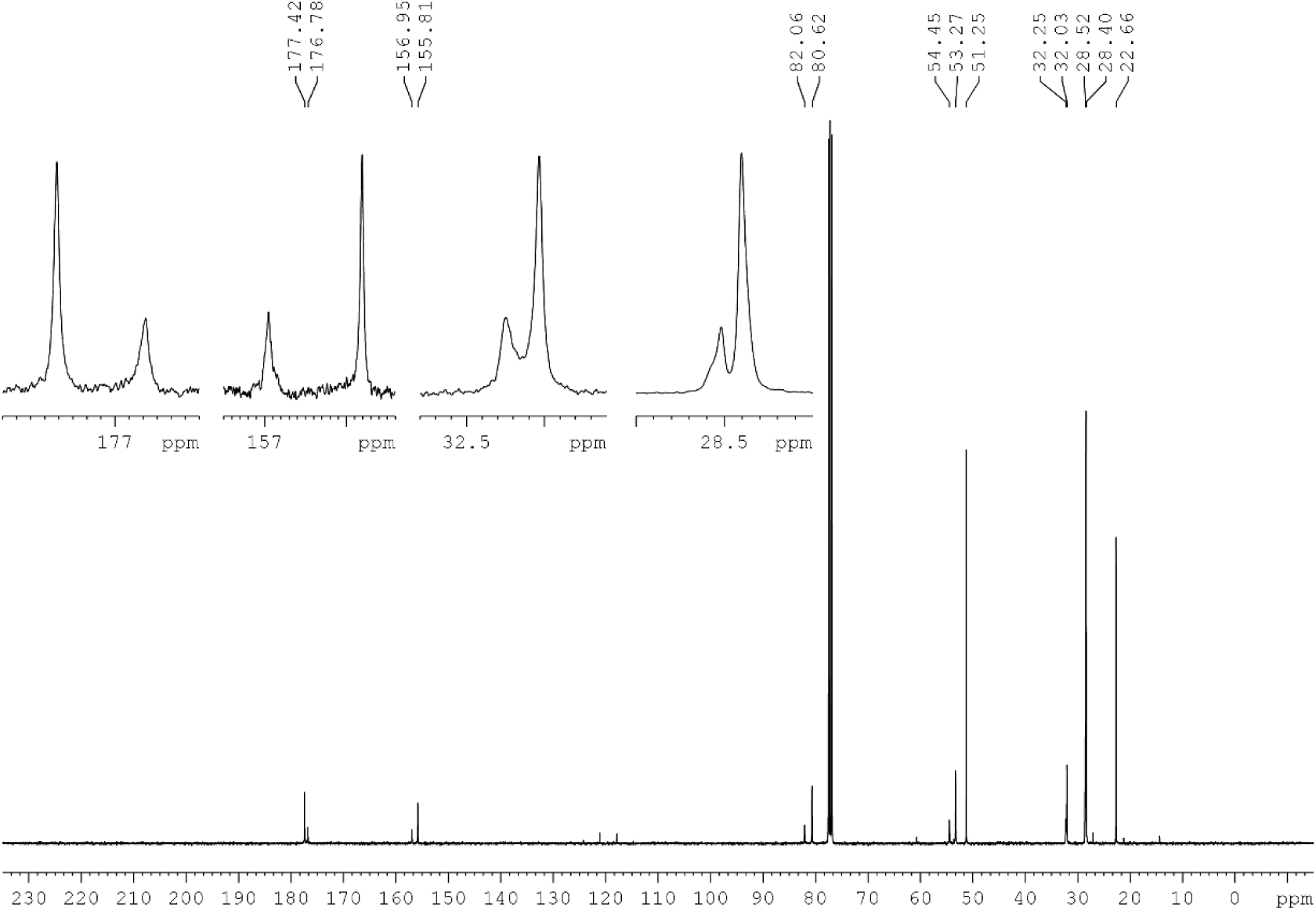
^13^C NMR spectrum of **2** (CDCl_3_, 100 MHz).

**Supplementary Figure S3.**
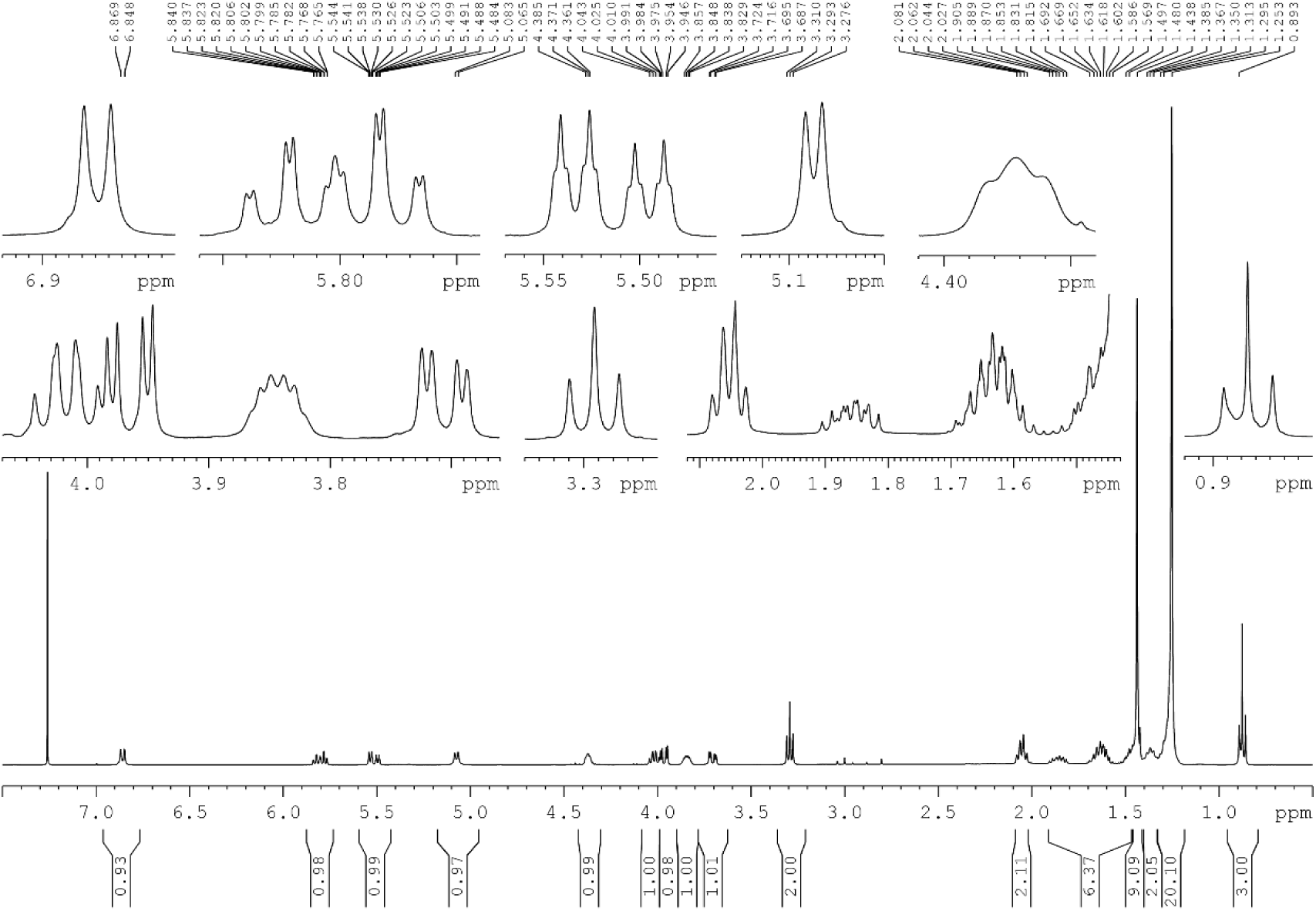
^1^H NMR spectrum of **3** (CDCl_3_, 400 MHz).

**Supplementary Figure S4.**
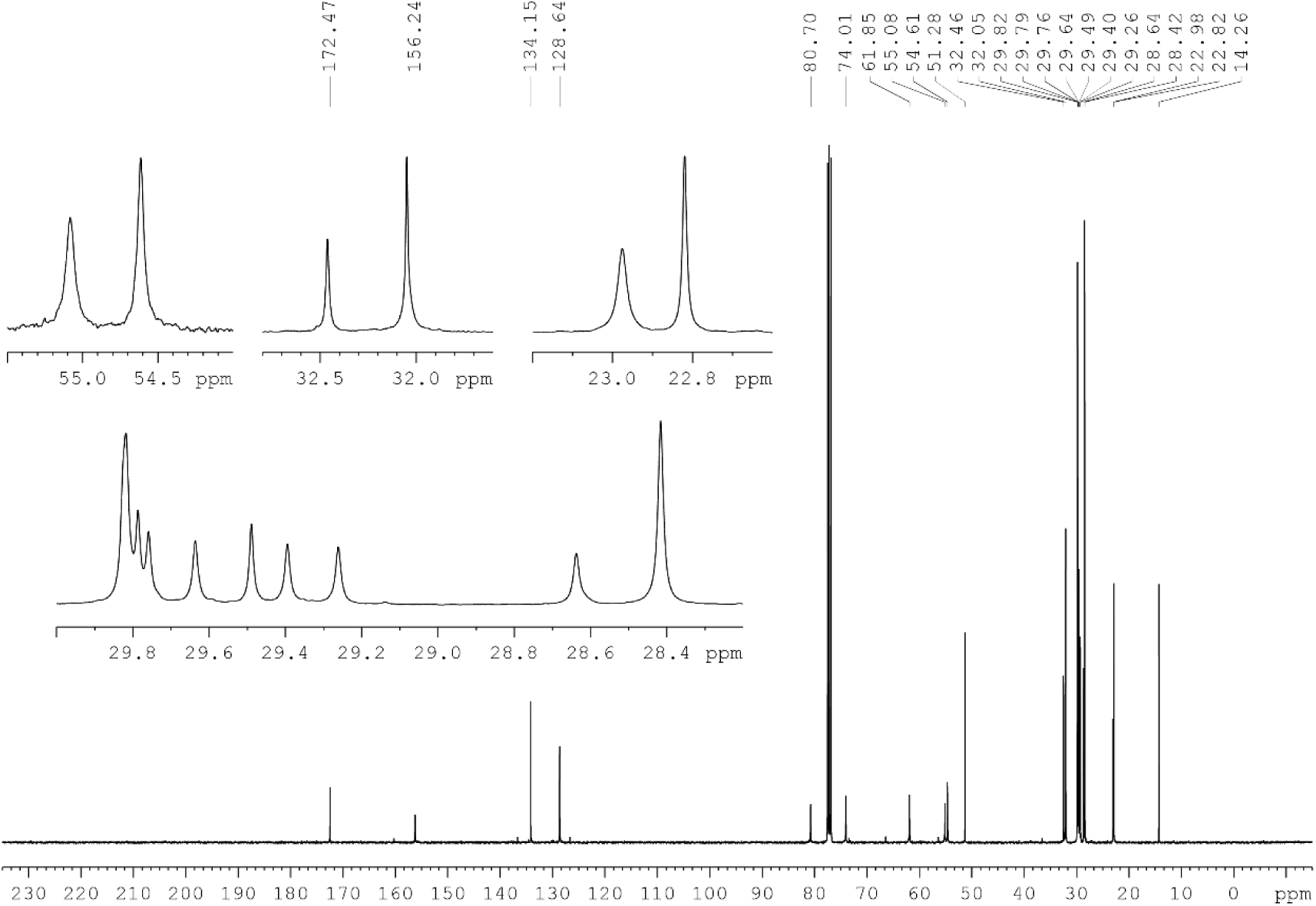
^13^C NMR spectrum of **3** (CDCl_3_, 100 MHz).

**Supplementary Figure S5.**
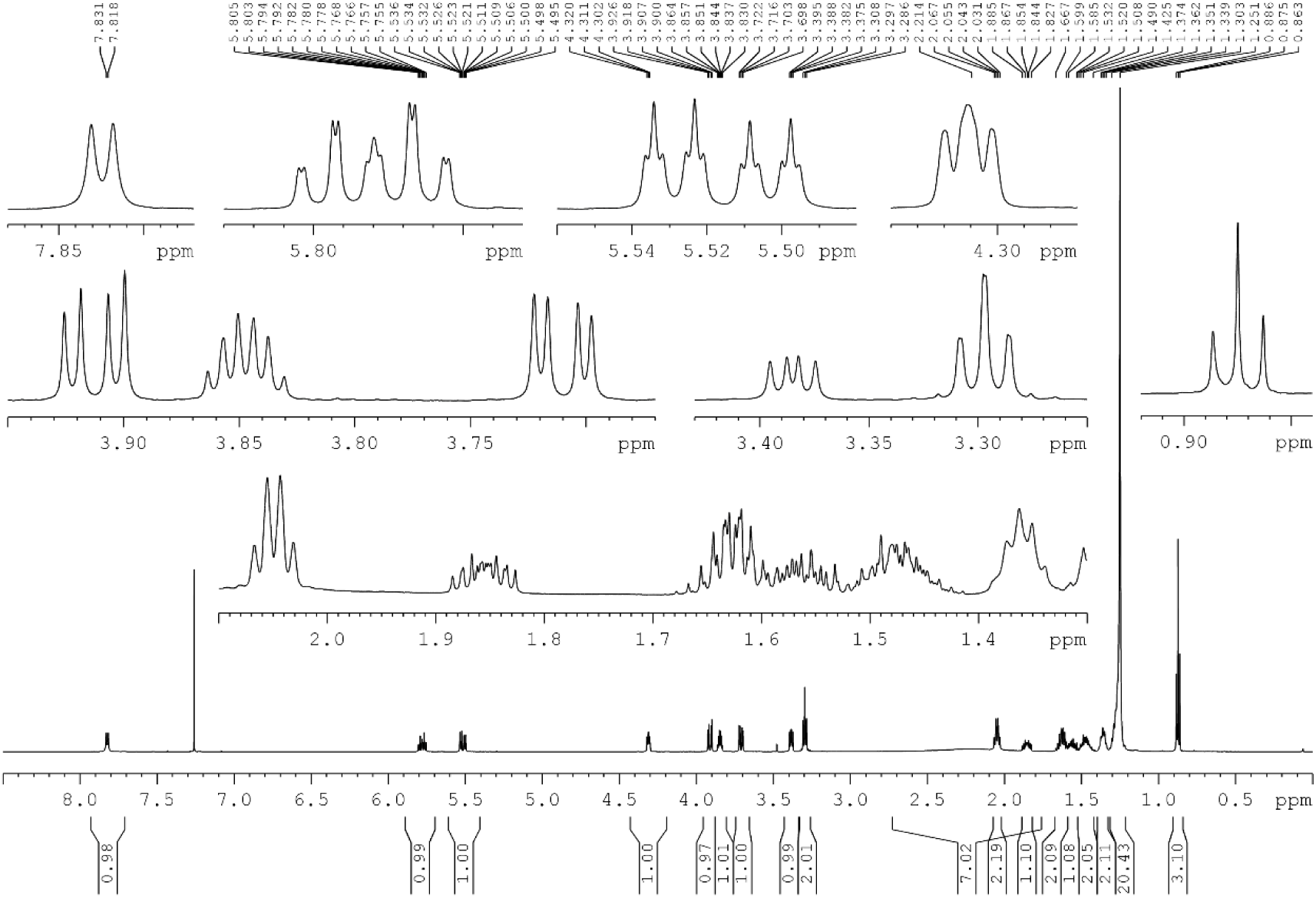
^1^H NMR spectrum of **4** (CDCl_3_, 600 MHz).

**Supplementary Figure S6.**
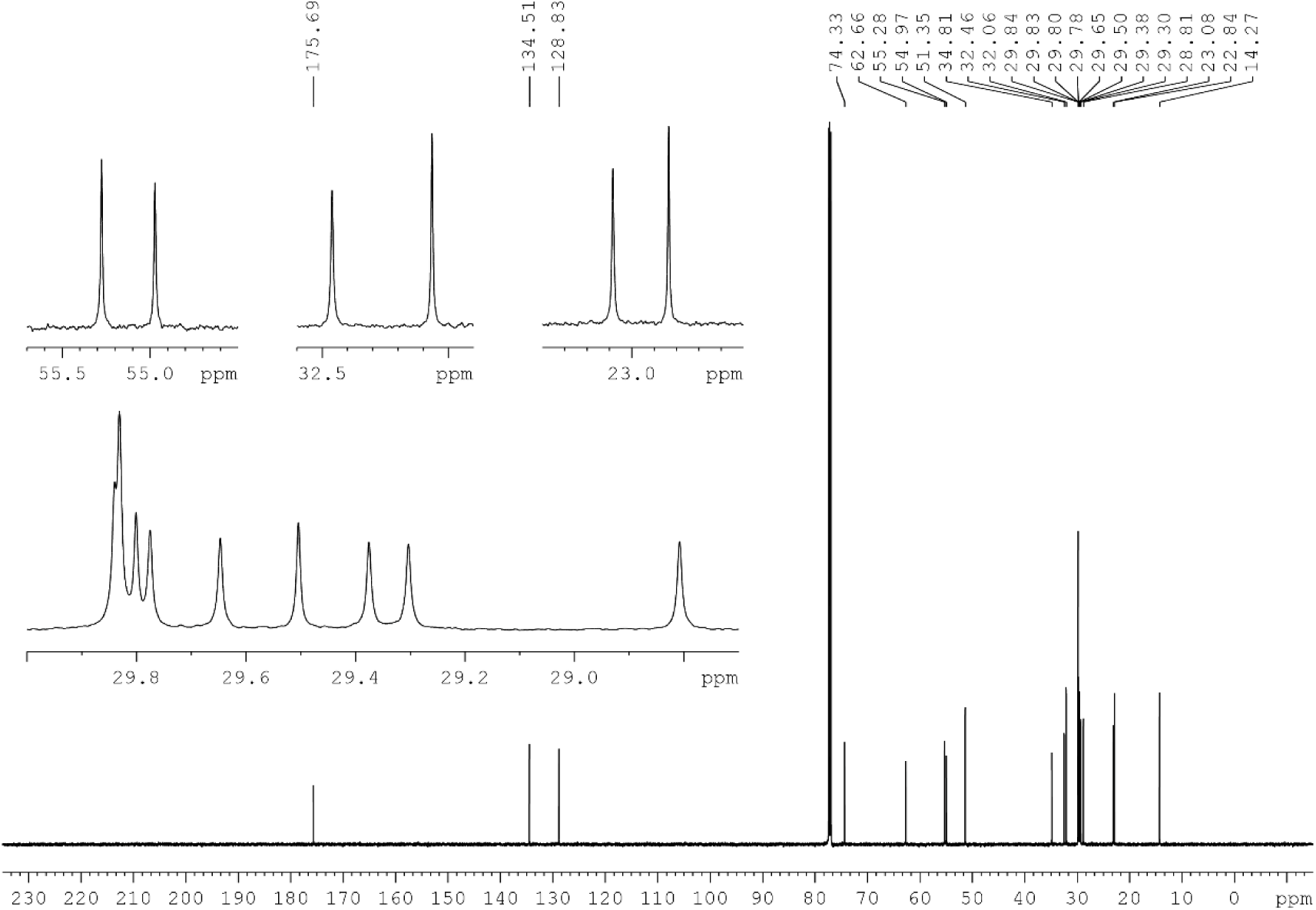
^13^C NMR spectrum of **4** (CDCl_3_, 150 MHz).

**Supplementary Figure S7.**
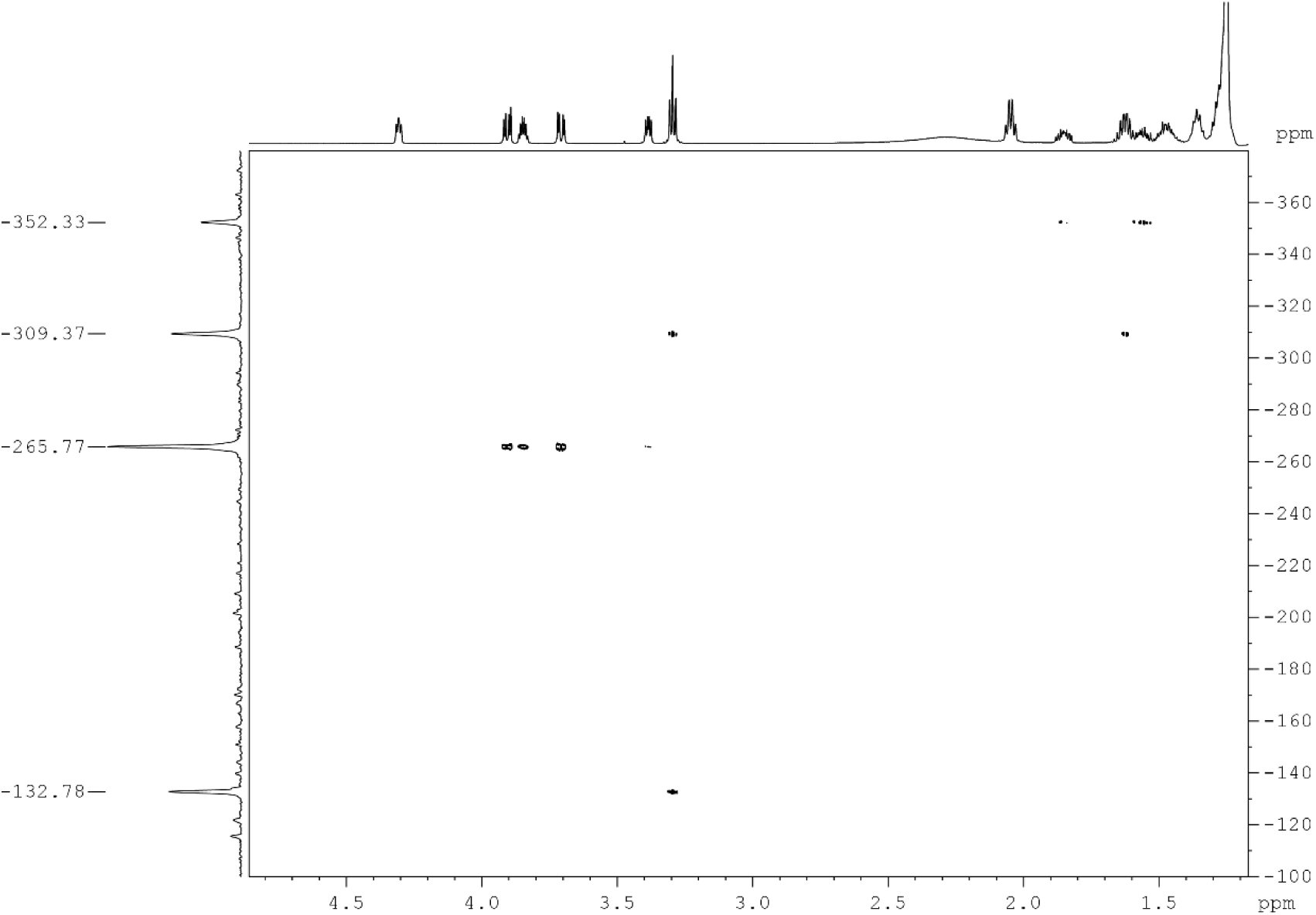
(^1^H,^15^N)-HMBC NMR spectrum of **4** (CDCl_3_, 600 MHz).

**Supplementary Figure S8.**
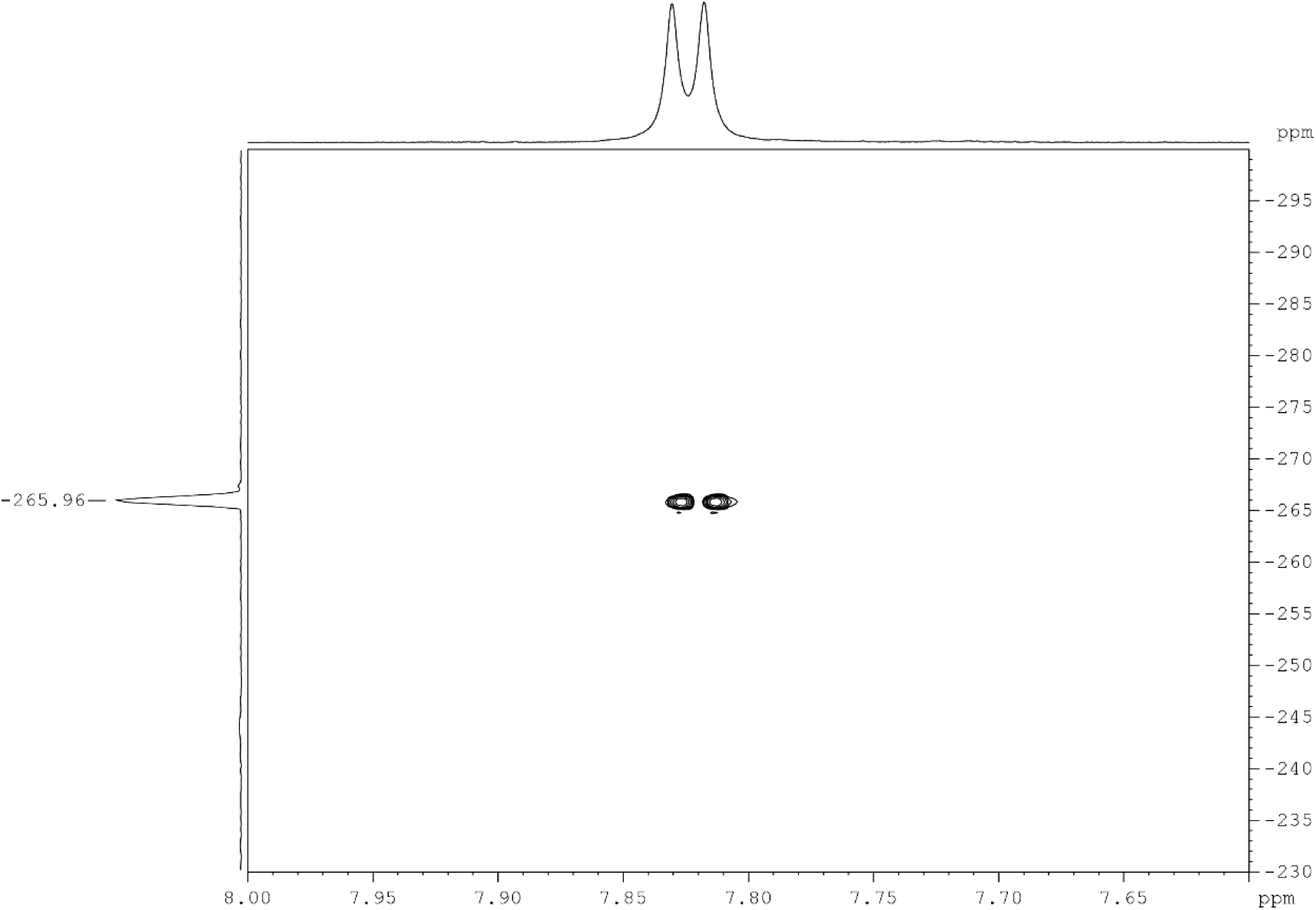
(^1^H,^15^N)-HSQC NMR spectrum of **4** (CDCl_3_, 600 MHz).

## Mass Spectra

**Supplementary Figure S9.**
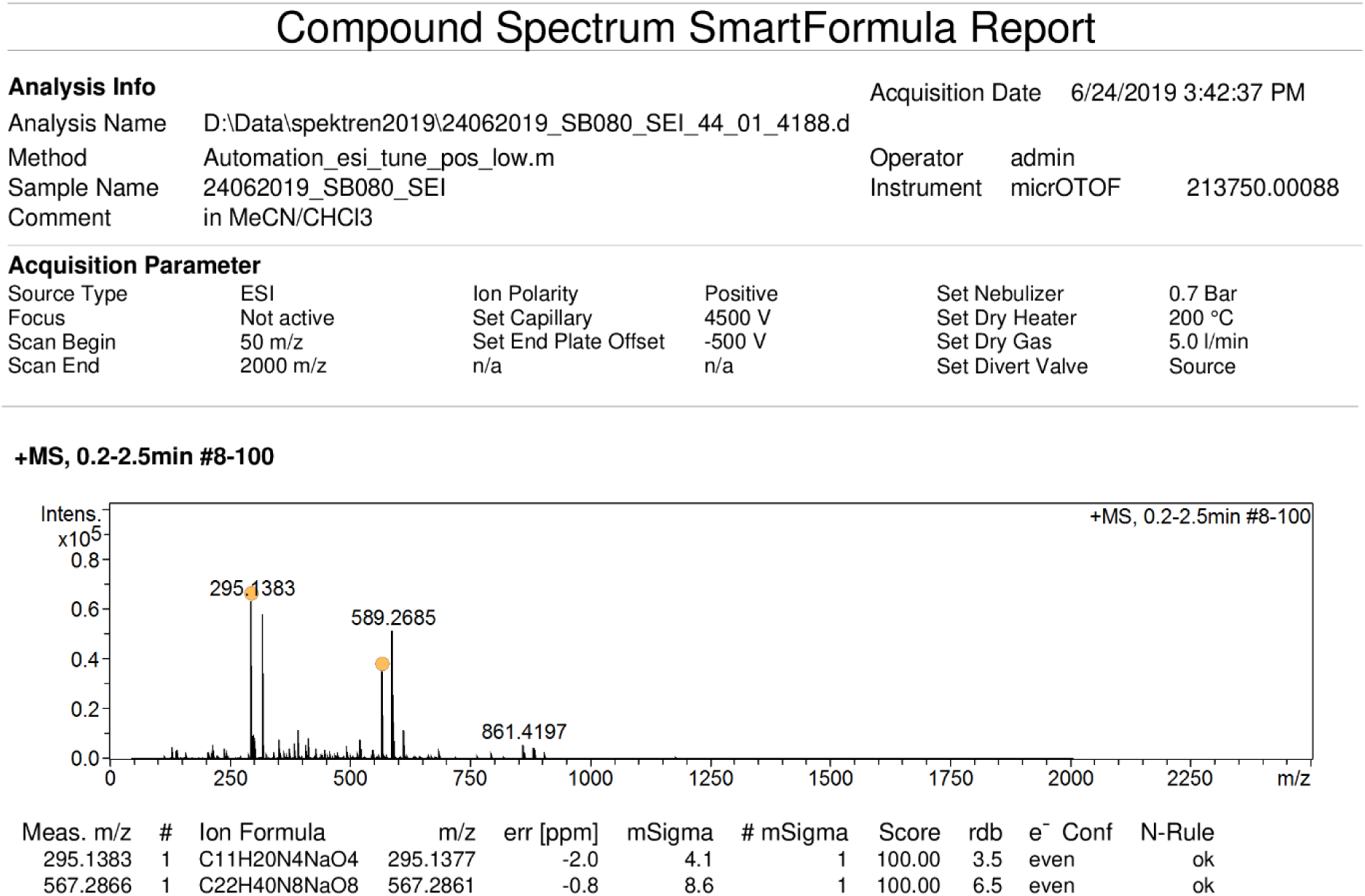
Mass spectrum of **2** (ESI^+^).

**Supplementary Figure S10.**
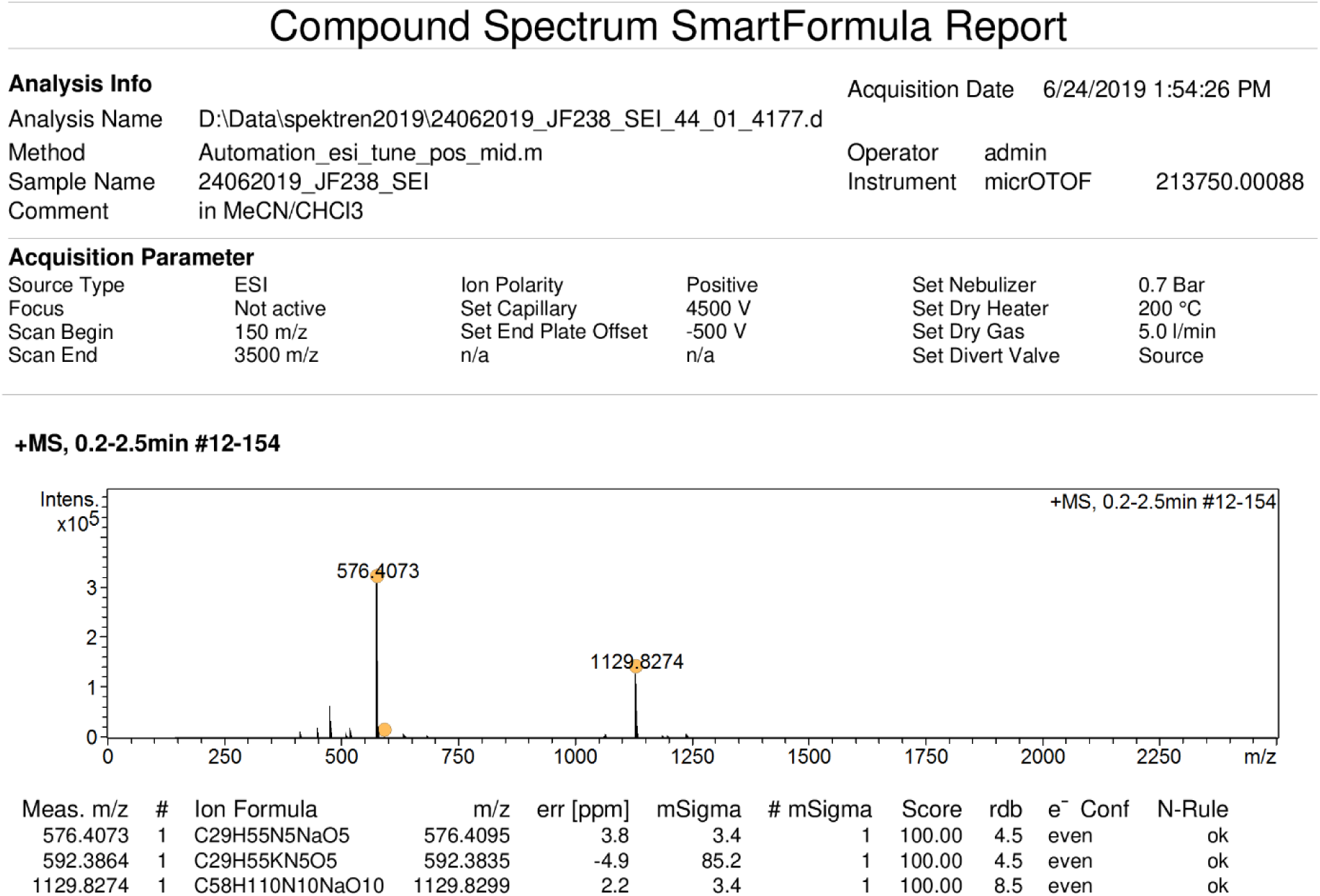
Mass spectrum of **3** (ESI^+^).

**Supplementary Figure S11.**
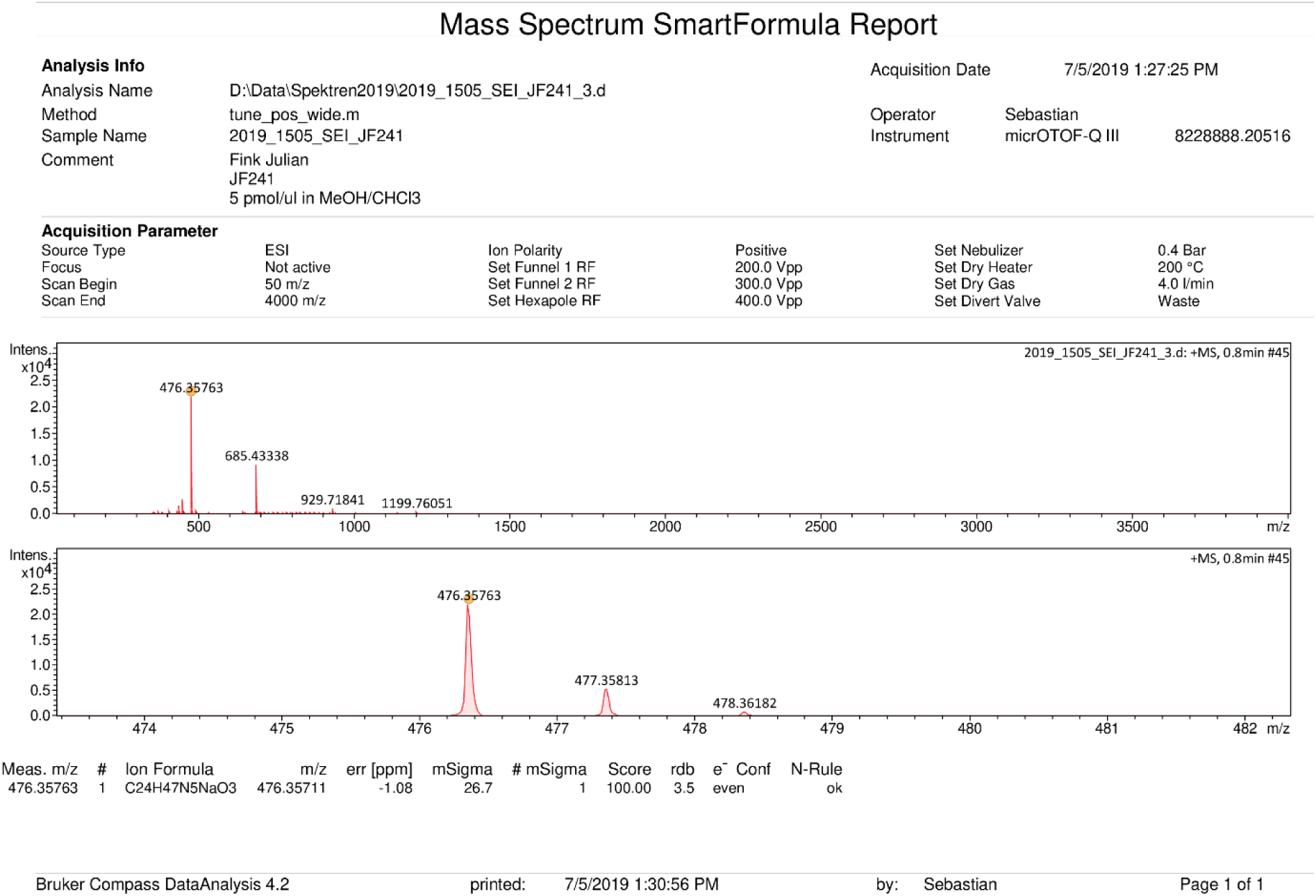
Mass spectrum of **4** (ESI^+^).

## IR Spectra

**Supplementary Figure S12.**
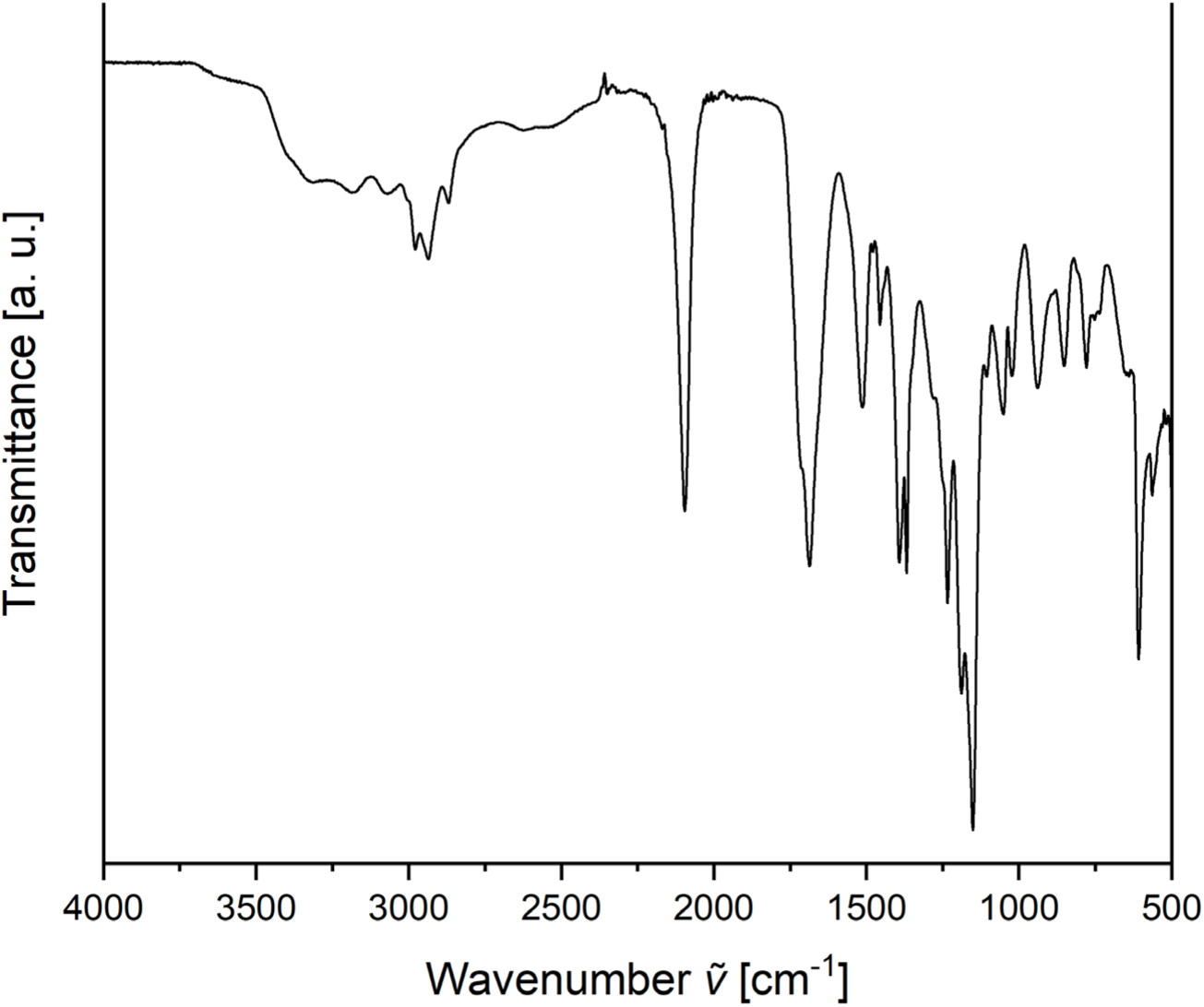
FTIR spectrum of **2** (ATR).

**Supplementary Figure S13.**
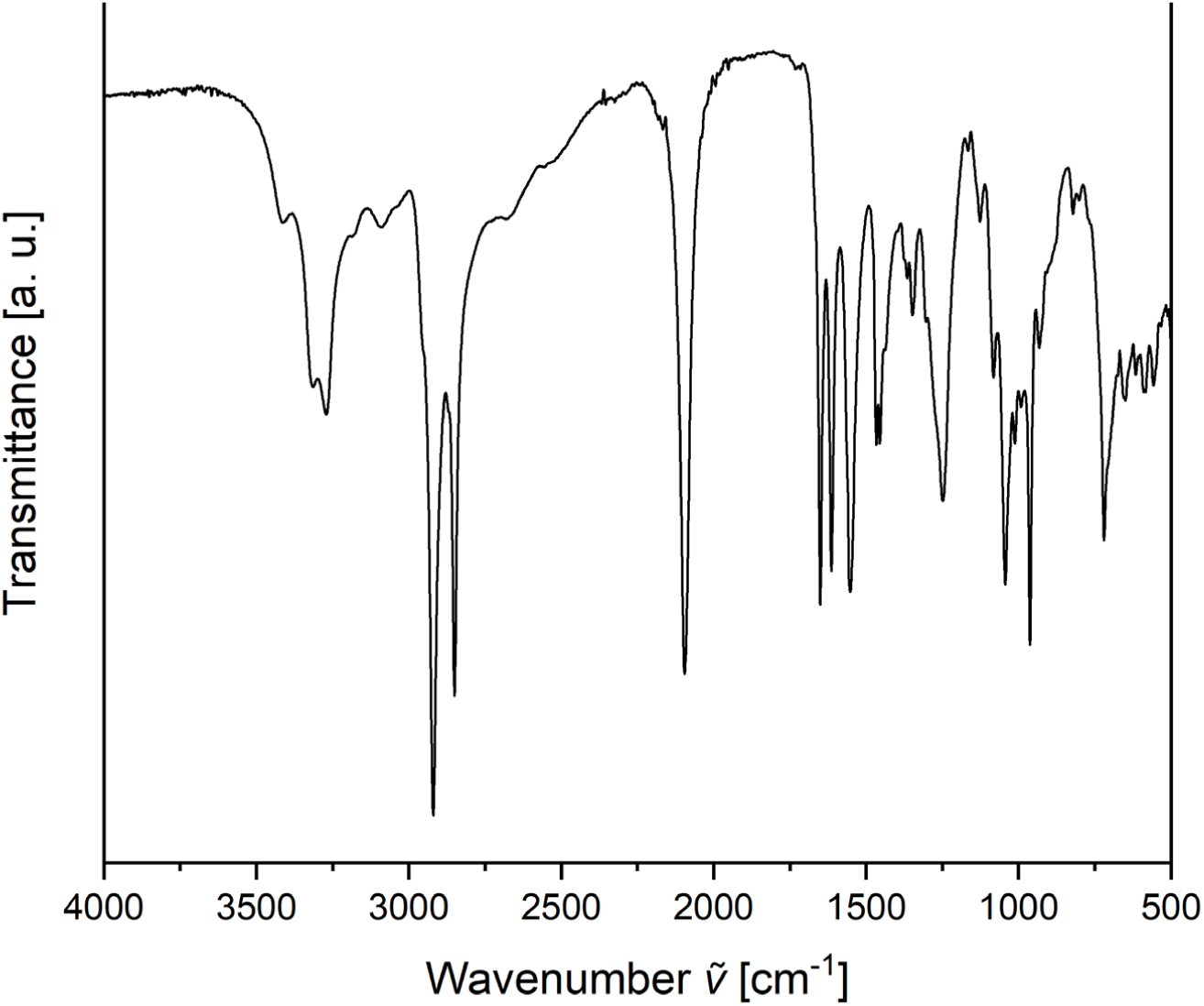
FTIR spectrum of **4** (ATR).

## ExM Images and Reference Experiments

**Supplementary Figure S14.**
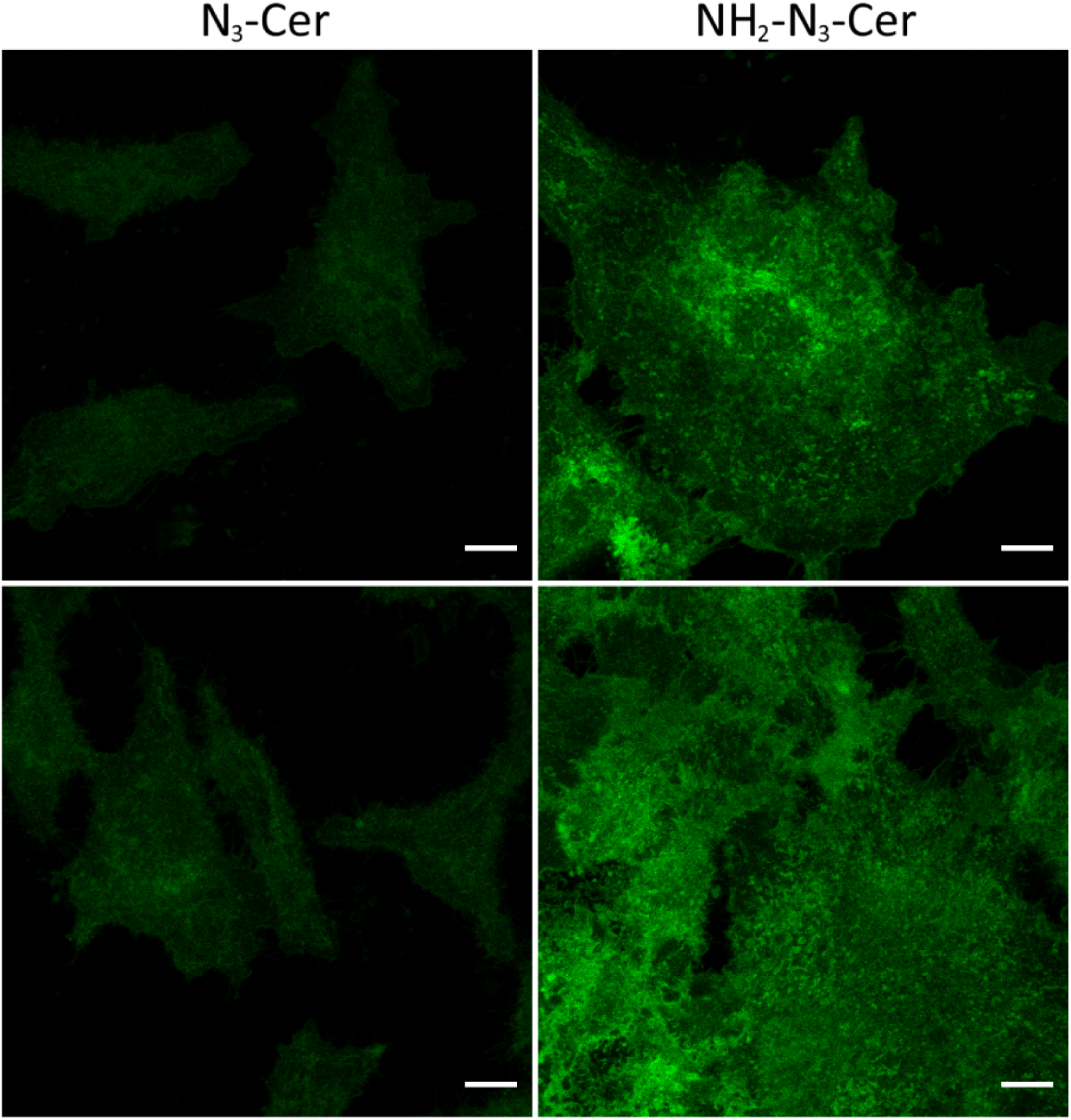
Ceramide signal after treatment with saponine. Hela229 cells were fed with ω-N_3_-C_6_-ceramide (N_3_-Cer) or α-NH_2_-ω-N_3_-C_6_-ceramide (NH_2_-N_3_-Cer) fixed, permeabilized with saponine and stained with DBCO-Alexa Fluor 488. Confocal fluorescence images show that N_3_-Cer is almost completely washed out while the NH_2_-N_3_-Cer signal remains preserved. Scale bars, 10 µm.

**Supplementary Figure S15.**
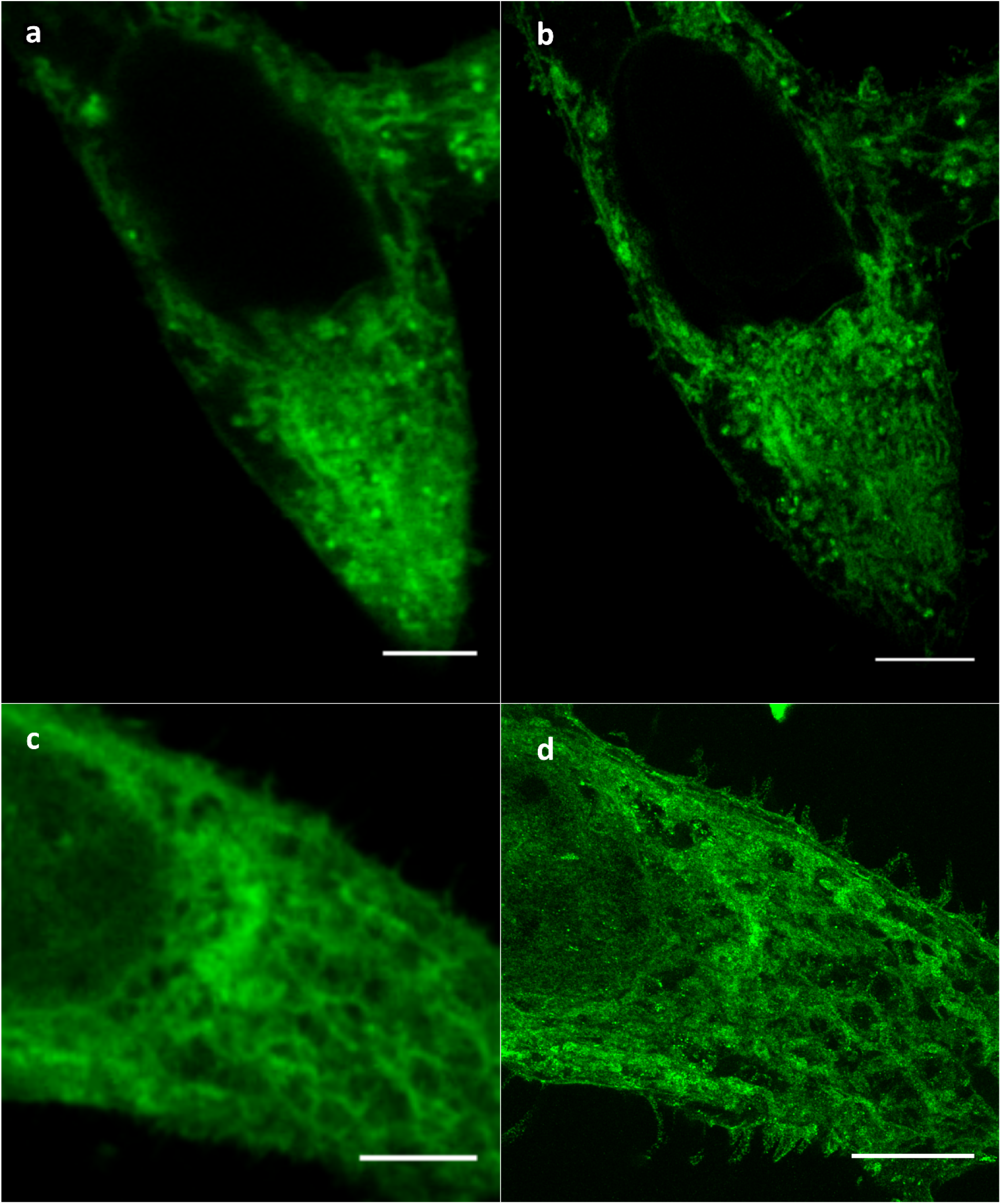
Confocal fluorescence images of the same HeLa229 cells pre- (a,c) and post-4x (b) and 10x (d) expansion. Cells were fed with α-NH_2_-ω-N_3_-C_6_-ceramide (NH_2_-N_3_-Cer), fixed, permeabilized, stained with DBCO-Alexa Fluor 488 and gelated. The images demonstrate isotropic expansion. The effective expansion factors were determined to 4.1x and 9.8x, respectively, from the cell’s diameters before and after expansion. Scale bars, unexpanded 5 µm (a,c), 4x expanded 20 µm (b) and 10x expanded 50 µm (d).

**Supplementary Figure S16.**
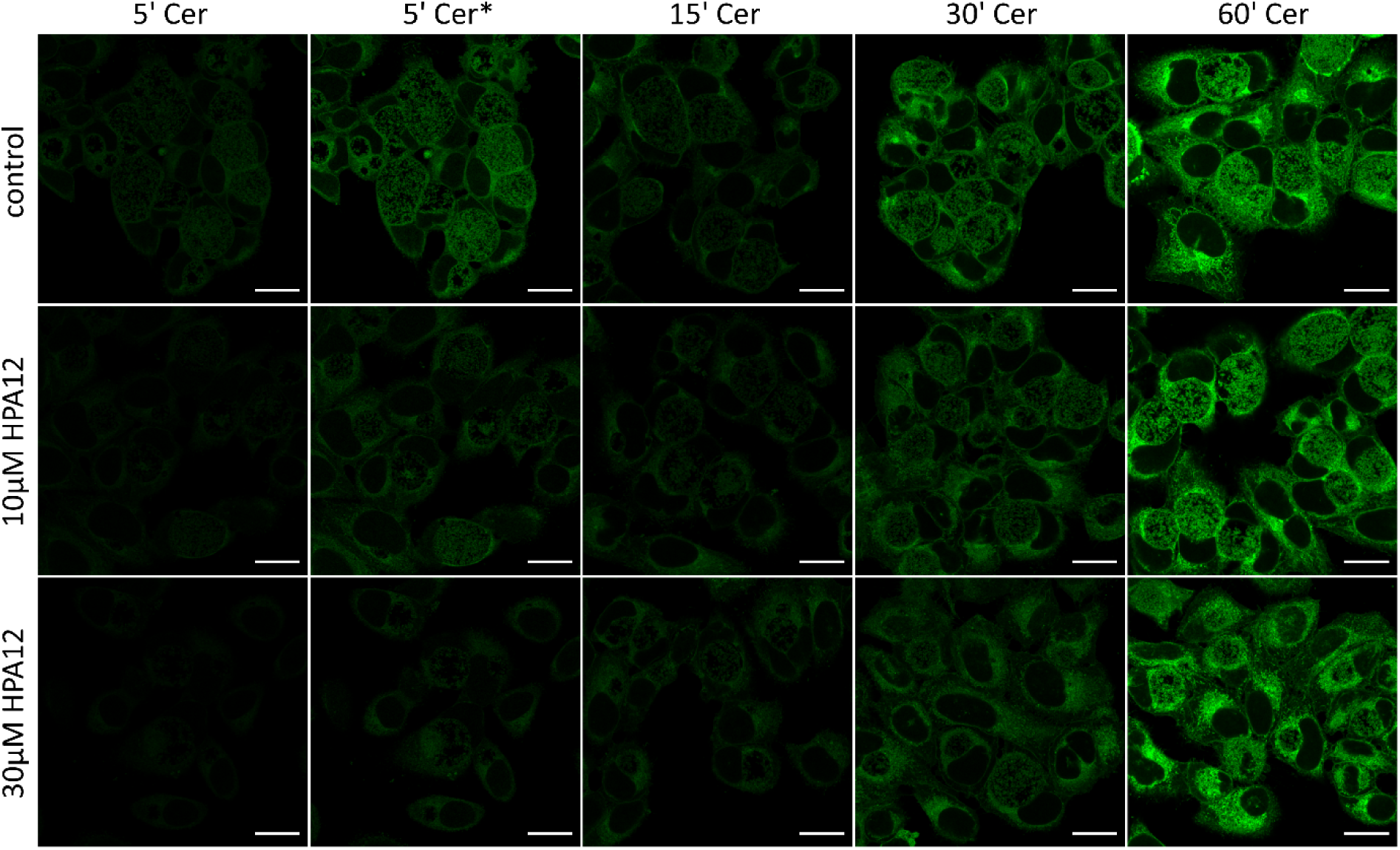
HPA-12 inhibits the uptake of α-NH_2_-ω-N_3_-C_6_-ceramide. HeLa229 cells were infected with *Chlamydia trachomatis* and treated with HPA-12 at 24 h of infection. 8 h later cells were fed with 10 µM α-NH_2_-ω-N_3_-C_6_-ceramide for 5-60 min, fixed, permeabilized and stained with DBCO-Alexa Fluor 488 (green). Confocal fluorescence images show that uptake of ceramides is reduced during the first 5 -15 min, while little to no difference is observed at longer incubation times. Scale bars, 20 µm.

**Supplementary Figure S17.**
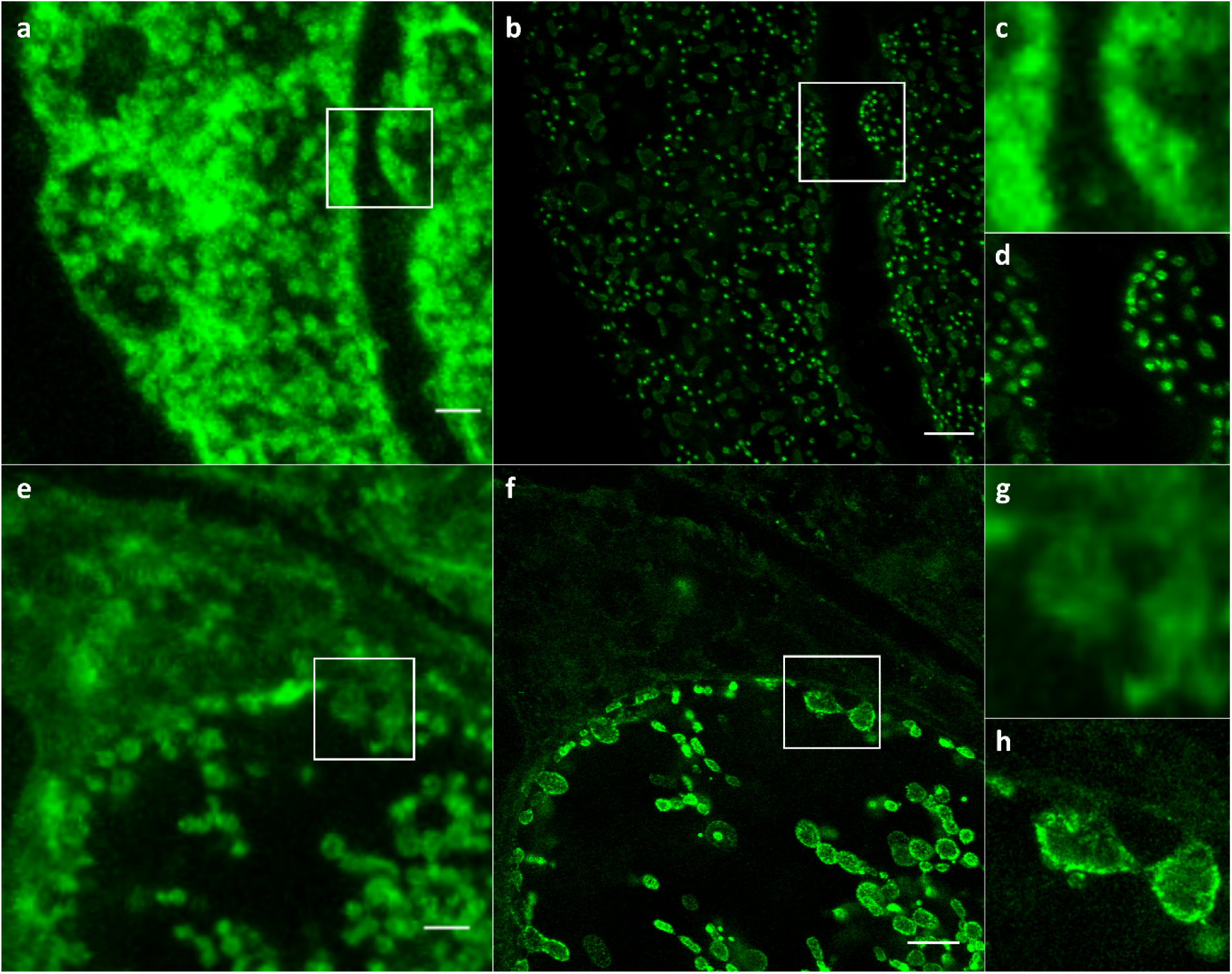
Comparing pre- and post-expanded infected cells. Confocal fluorescence images of the same areas of HeLa229 cells infected with *S. negevensis* (a-d) or *C. trachomatis* (e-h), before (a,c,e,g) and after (b,d,f,h) 10x expansion. The infected cells were fed α-NH_2_-ω-N_3_-C_6_-ceramide, fixed, permeabilized, and stained with DBCO-Alexa 488 following gelation. (c,d) and (g,h) show magnified views of the regions outlined by the white boxes in the main images. Scale bars, unexpanded 2 µm and 10x expanded 20 µm.

**Supplementary Figure S18.**
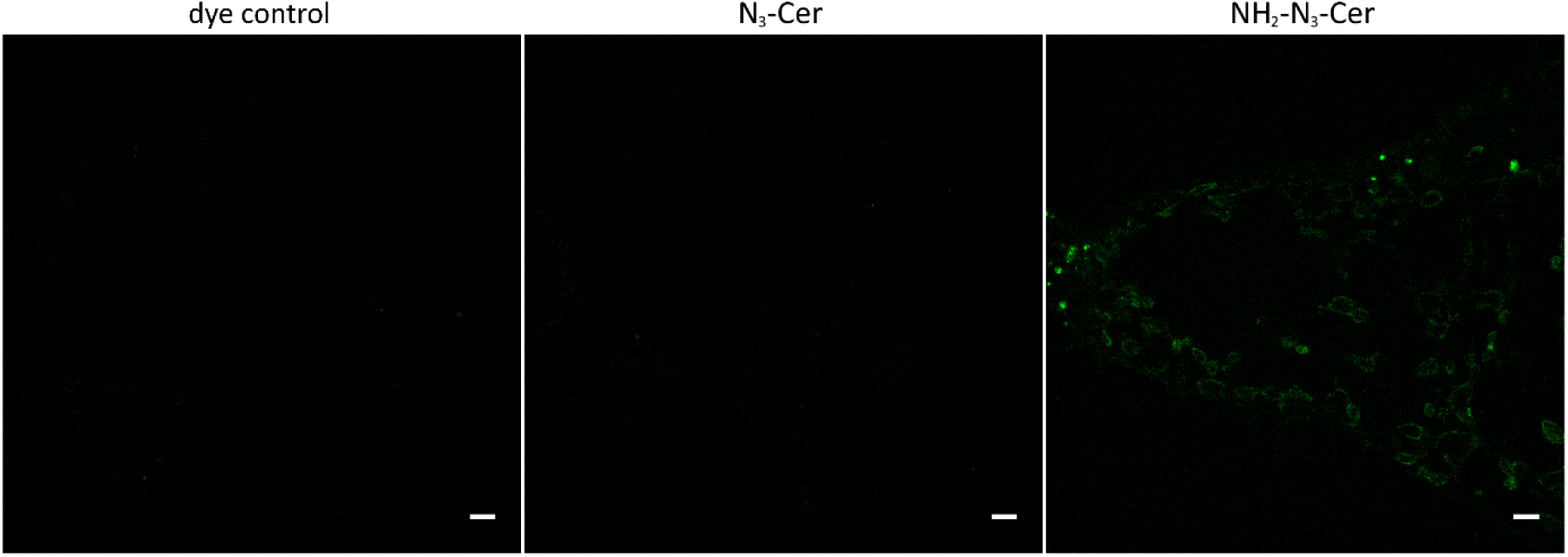
Confocal fluorescence images of HeLa229 cells infected with *Chlamydia trachmomatis* for 24 h, without ceramide feeding (dye control), with ω-N_3_-C_6_-ceramide (N_3_-Cer), or with α-NH_2_-ω-N_3_-C_6_-ceramide (NH_2_-N_3_-Cer), fixed, permeabilized and stained with DBCO-Alexa Fluor 488. Only after feeding with α-NH_2_-ω-N_3_-C_6_-ceramide (NH_2_-N_3_-Cer) the bacteria are visualized. Scale bars,10 µm.

**Supplementary Figure S19.**
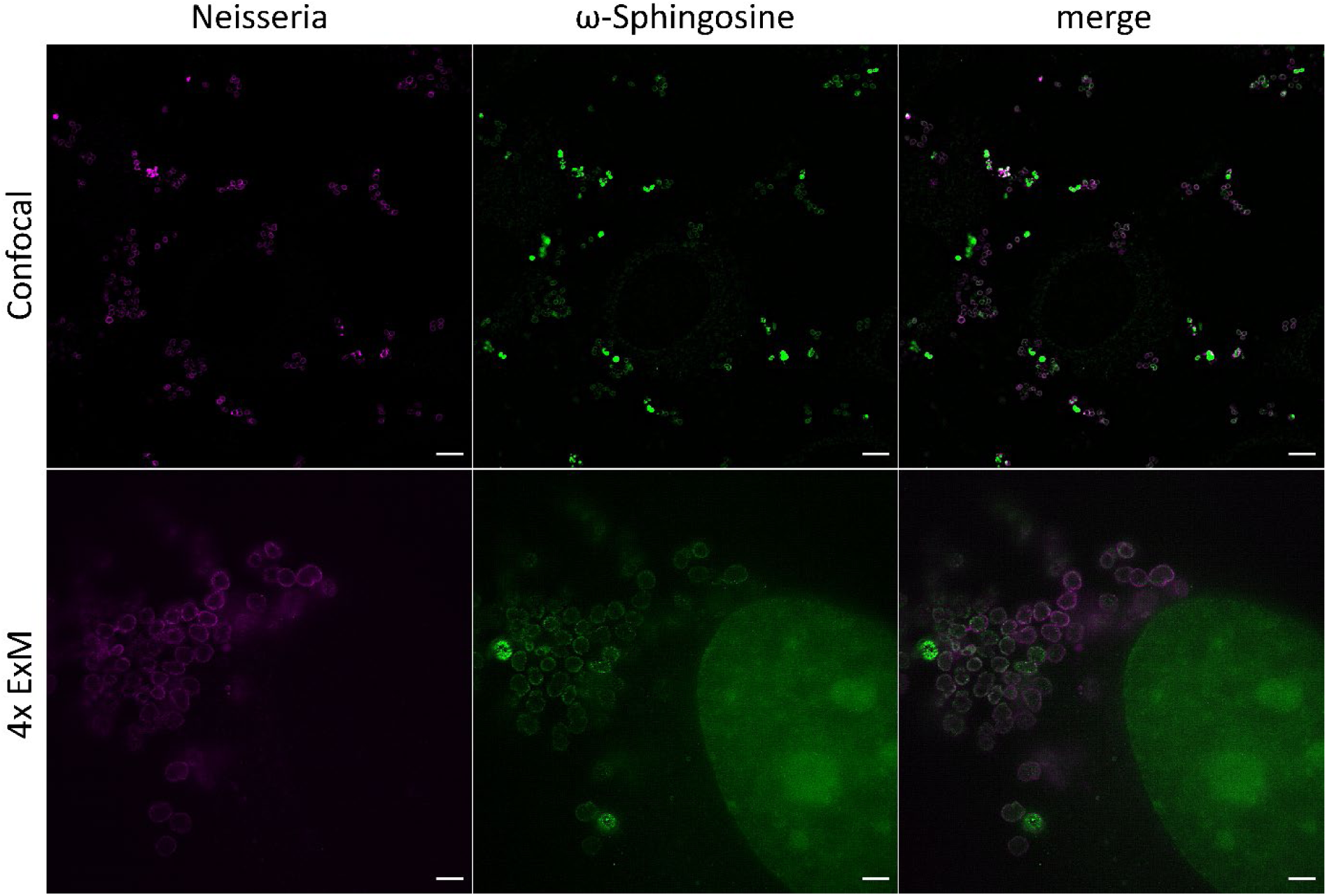
Confocal (upper row) and 4x ExM-SIM images (lower row) of Hela229 infected cells infected with *Neisseria gonorrhoeae* for 4 h, fed with ω-sphingosine, fixed, permeabilized and stained with DIBO-Alexa Flour 488 (green) and anti-Neisseria (magenta). SIM images clearly show incorporation of ω-sphingosine into the membrane of Neisseria. Scale bars, 5 µm (unexpanded confocal), 4 µm (expanded SIM).

**Supplementary Figure S20.**
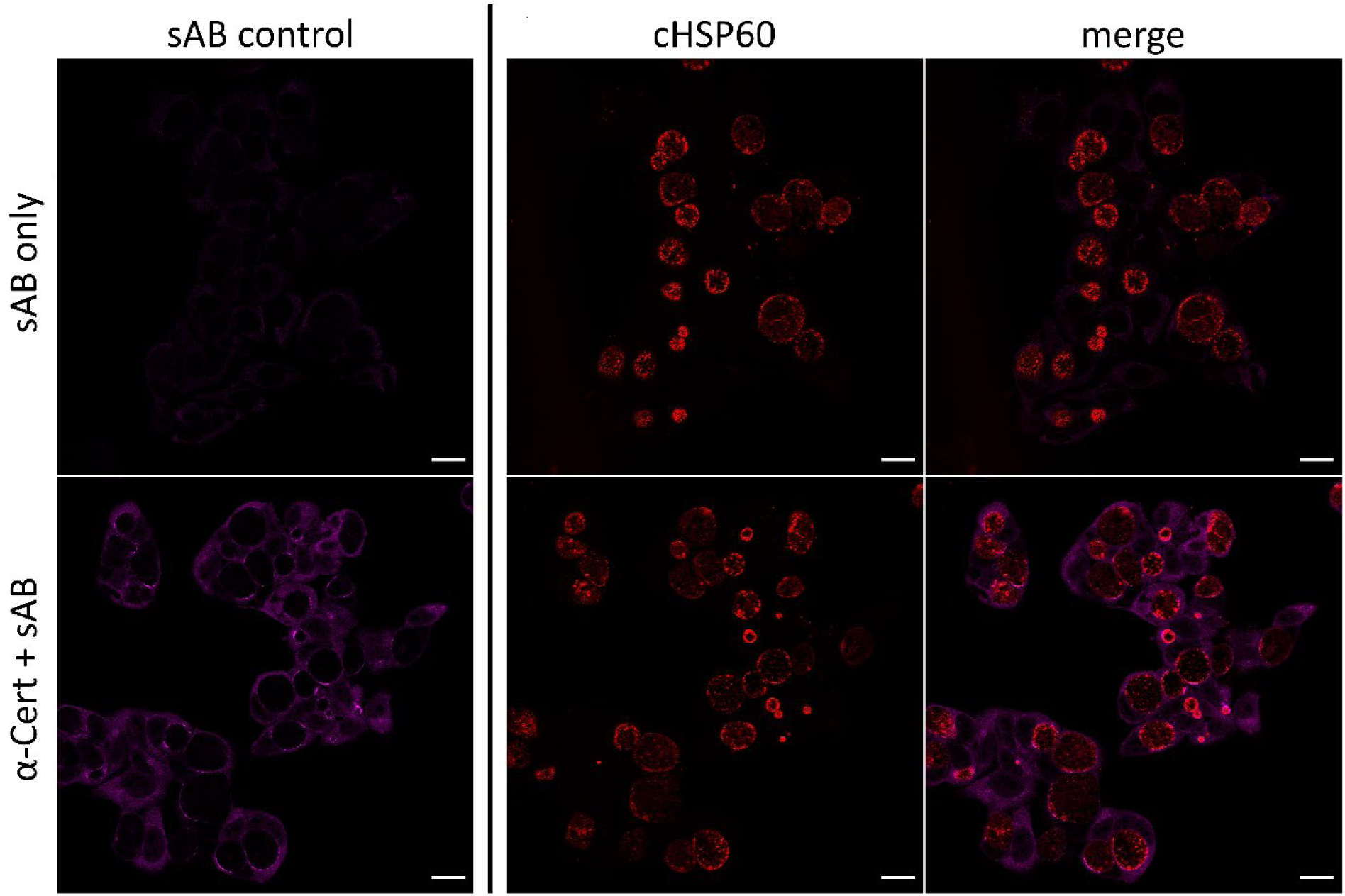
Confocal images of unexpanded infected Hela229 cells infected with *Chlamydia trachomatis* for 24 h, fed with α-NH_2_-ω-N_3_-C_6_-ceramide, fixed, permeabilized and immunolabeled for α-Cert with ATTO 647N (magenta) and chlamydial HSP60 with Cy3 (red). Labeling with secondary antibodies alone does not show any staining of chlamydial particles (sAB only/sAB control). Scale bars, 20 µm.

**Supplementary Figure S21.**
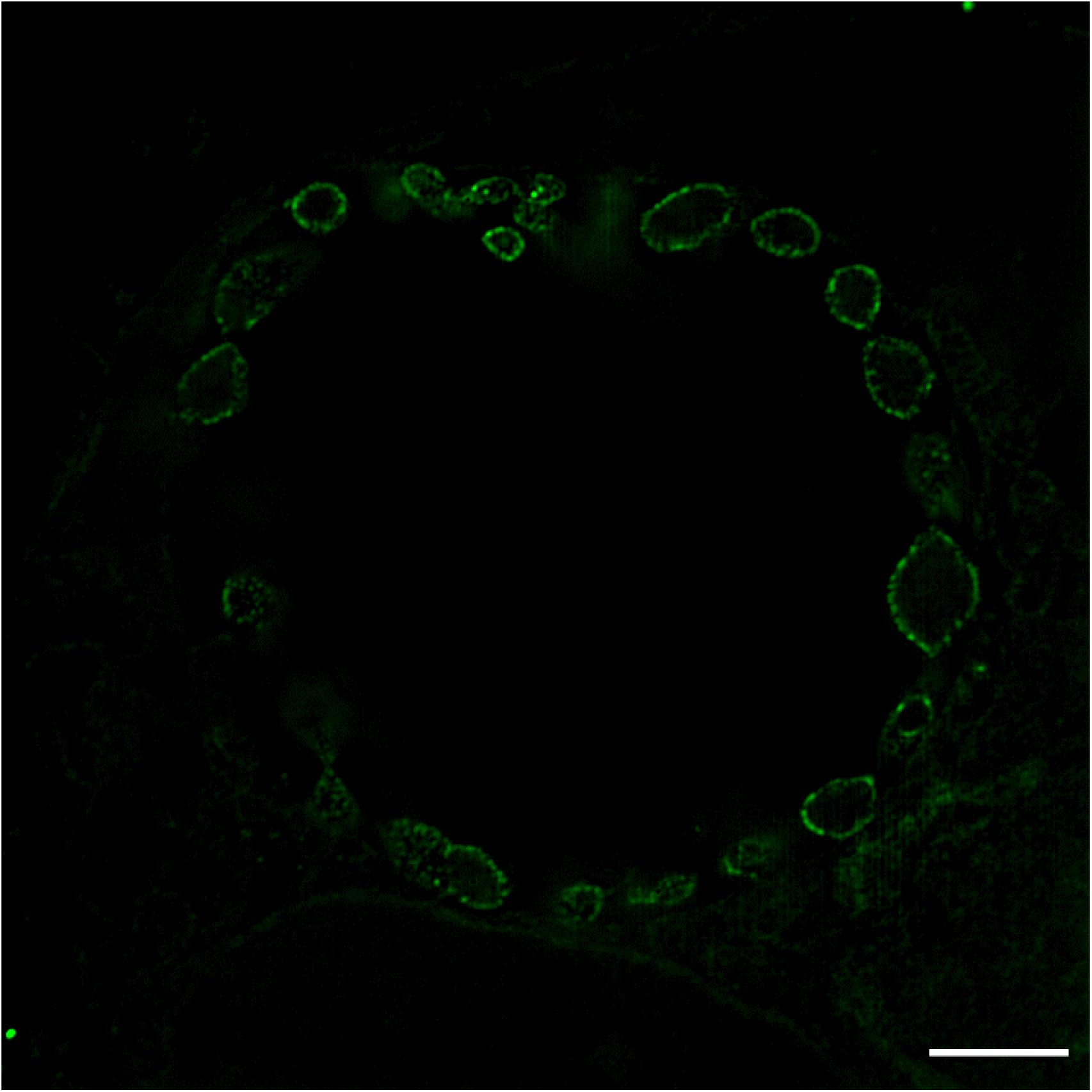
10x ExM-SIM image of Hela229 cells infected with *Chlamydia trachomatis* for 24 h, fed with α-NH_2_-ω-N_3_-C_6_-ceramide, fixed, permeabilized and stained with DBCO-Alexa Fluor 488. Chlamydia are clearly located at the inclusion membrane. Scale bar, 10 µm.

**Supplementary Figure S22.**
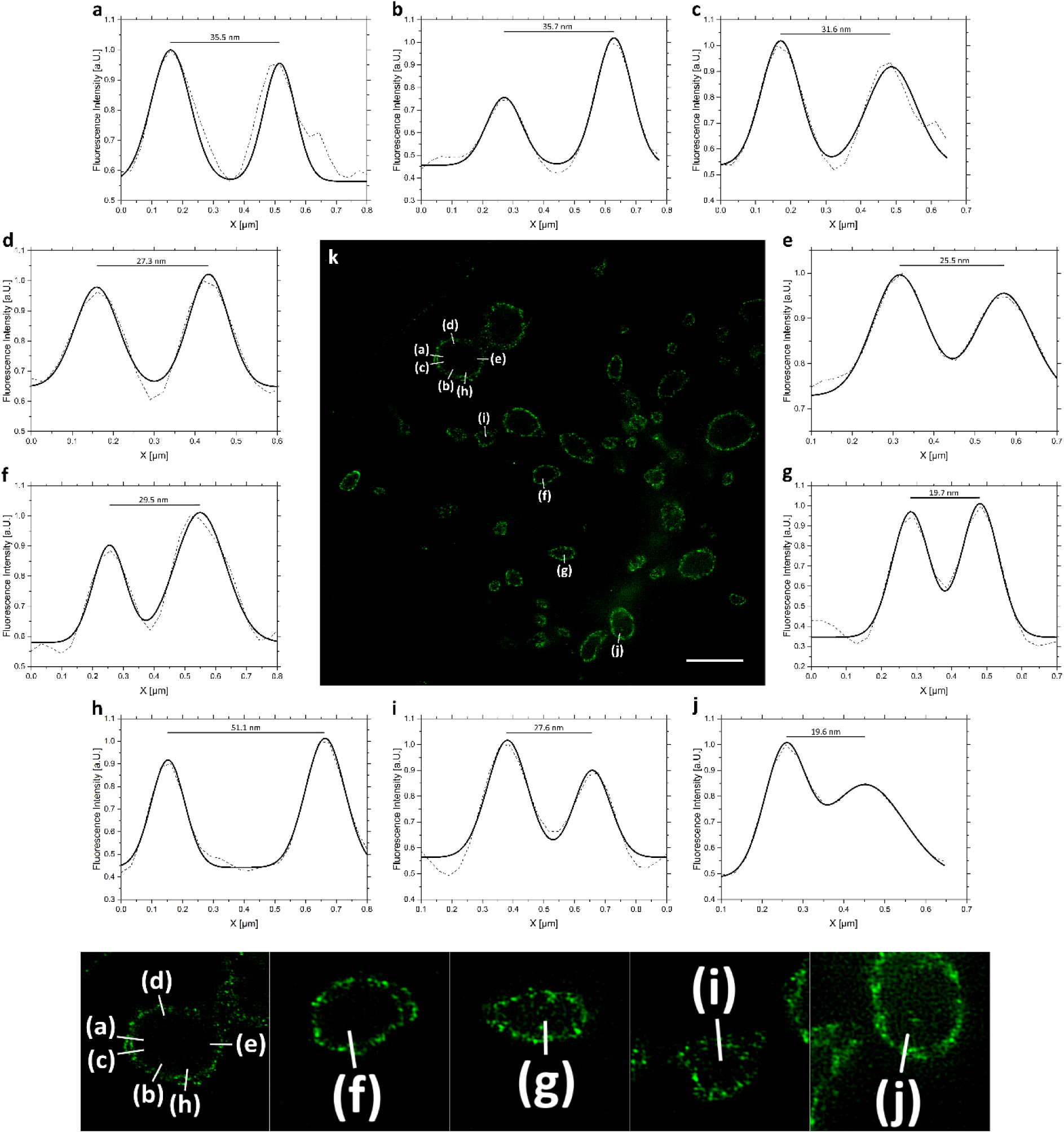
Cross sectional intensity profiles (**a**-**j**) of 10x expanded *C. trachomatis* fed with α-NH_2_-ω-N_3_-C_6_-Ceramide (green) in Hela229 cells. The distance between the inner and outer membrane varies between 19.7 nm and 51.1 nm. (k) SIM image of one of three 10x expanded sample used for data analysis.

**Supplementary Figure S23.**
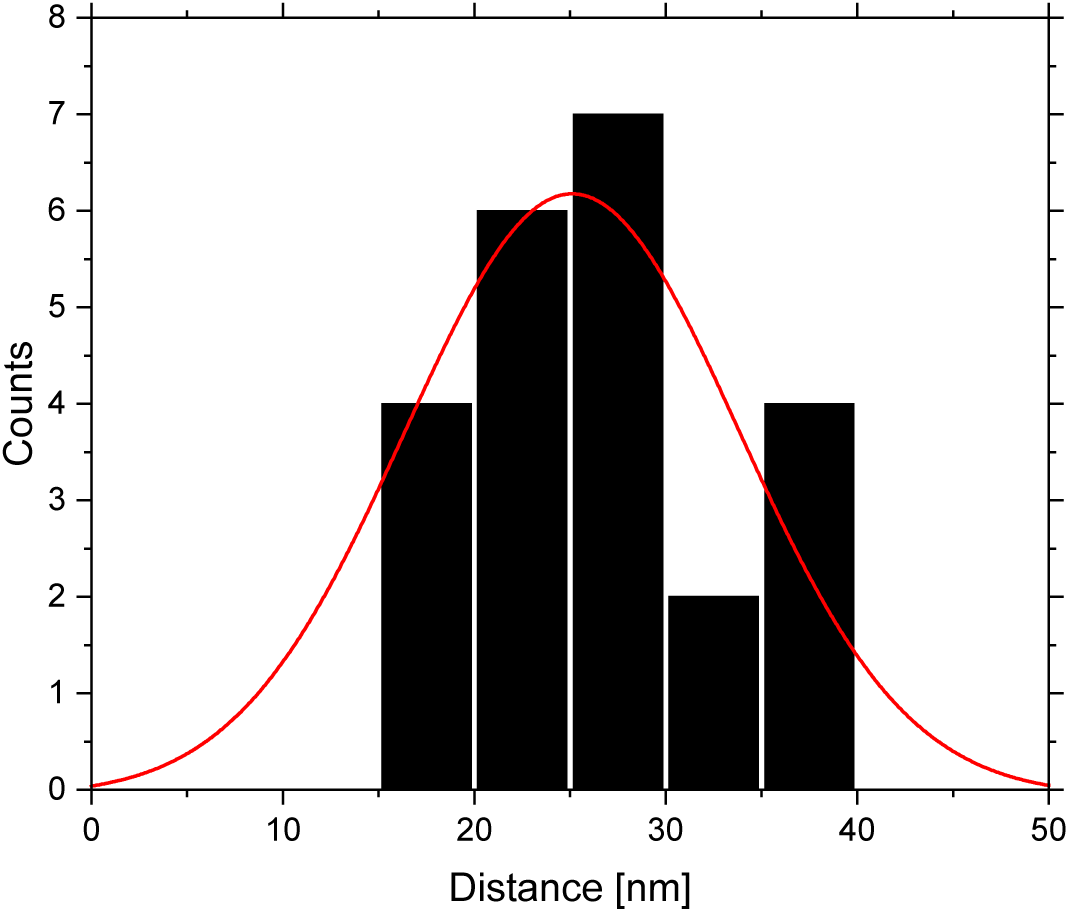
Histogram and fit of 23 distances determined from cross sectional intensity profiles from three biological replicates infected with *C. trachomatis* and fed with α-NH_2_-ω-N_3_-C_6_-Ceramide (Supplementary Figure S22). Only those bacteria were selected whose orientation allowed us to visualize spatially separated OM and IM (i.e. frontal views of bacteria) and determine the distance between the two membranes to 27.6 ± 7.7 nm (s.d.).

**Supplementary Video 1.** 10x ExM SIM z-stack of Hela229 cells infected with *Chlamydia trachomatis* for 24 h, fed with α-NH_2_-ω-N_3_-C_6_-ceramide, fixed, permeabilized and stained with DBCO-Alexa Fluor 488. Chlamydia are clearly located at the inclusion membrane. Scale bar, 10 µm.

